# Moderate DNA methylation changes associated with nitrogen remobilization and leaf senescence in *Arabidopsis*

**DOI:** 10.1101/2021.09.17.460744

**Authors:** Emil Vatov, Ulrike Zentgraf, Uwe Ludewig

## Abstract

- The lifespan of plants and tissues is restricted by environmental and genetic components. Following the transition to reproductive growth, leaf senescence ceases cellular life in monocarpic plants to remobilize nutrients to storage organs.
- We observed altered leaf to seed ratios, faster senescence progression and enhanced nitrogen remobilization from the leaves in two methylation mutants (*ros1* and the triple *dmr1/2 cmt3* knockout).
- DNA methylation in wild type Col-0 leaves initially moderately declined with progressing leaf senescence, predominantly in the CG context, while the ultimate phase of leaf discoloration was associated with moderate *de novo* methylation of cytosines, primarily in the CHH context.
- Relatively few differentially methylated regions, including one in the *ROS1* promoter linked to the down-regulation of *ROS1,* were present, but these were unrelated to known senescence-associated genes.
- Differential methylation patterns were identified in transcription factor binding sites, such as the W-boxes that are targeted by WRKYs, which impaired transcription factor binding when methylated *in vitro*.
- Mutants that are defective in DNA methylation showed distinct nitrogen remobilization, which was associated with altered patterns of leaf senescence progression. But moderate methylome changes during leaf senescence were not specifically associated with up-regulated genes during senescence.

## Introduction

Senescence in monocarpic plants, such as *Arabidopsis*, is a developmentally regulated process that leads to the controlled death of the whole organism, except for the dormant seeds. After the transition to reproductive growth, not only the fully expanded leaves, but all leaves undergo senescence processes and are turned into source leaves for the newly developing flowers and seeds. The systematic dismantling of the chloroplasts is characteristic for leaf senescence, resulting in a visible loss of green coloration. RuBisCo (ribulose-1,5-bisphosphate carboxylase-oxygenase), which is involved in photosynthesis and locates in the chloroplast stroma, is the most abundant protein in leaf tissue and, therefore, accounts for a large proportion of the nitrogen within a leaf. During senescence, nitrogen (N) remobilization is thus coupled to chloroplast dismantling; the degradation of photosynthesis-related proteins results in reduced photosynthetic capacity (Diaz et al., 2008; Masclaux-Daubresse et al., 2010; Havé et al., 2017). Even though this compromises carbohydrate assimilation, soluble sugars and hexoses with signalling roles transiently increase in leaves during senescence (Wingler et al., 2006).

In addition to their deteriorative power, reactive oxygen species (ROS and, in particular, H_2_O_2_) increase during the transition to reproductive growth and, in turn, correlate with the expression of many senescence-associated genes (SAGs) including transcriptional regulators. The main transcription factor families orchestrating the process are NAC, WRKY, C2H2 and MYB (Breeze et al., 2011; Liebsch and Keech, 2016). WRKY53 appears to play a key role in early senescence regulation and is tightly controlled by many different mechanisms at the transcriptional and posttranscriptional level (Zentgraf and Doll, 2019). As almost all WRKY factors carry DNA-binding sites for WRKY factors in their promoters (Dong et al., 2003), WRKY factors regulate each other in complex regulatory networks. For example, WRKY18 has been characterized as an upstream negative regulator, downstream target and protein interaction partner of WRKY53. Moreover, WKRY25 influences the expression of *WRKY53* and *WRKY18* in a redox-dependent manner but is itself also involved in regulating intracellular hydrogen peroxide concentrations (Doll et al., 2020). Moreover, WRKY53 and WRKY18 have been identified as regulators of the sugar response gene *GPT2* via the recruitment of HAC1 (Chen et al., 2019).

The involvement of epigenetic mechanisms, in the form of active histone modifications related to chromatin decondensation and the regulation of key senescence regulators, e.g. *WRKY53*, has also been observed (Ay et al., 2009; Brusslan et al., 2012). The expression of the *WRKY53* locus itself is under epigenetic control and the WRKY53 protein participates in the epigenetic control of other senescence regulators, i.e. the SANT-domain-containing protein POWERDRESS recruits the histone deacetylase HDA9 to the W-box-containing promoter regions of negative senescence regulator genes (e.g. *WRKY57, APG9,* and *NPX1*) with the help of WRKY53 (Chen et al., 2016).

Whereas ageing in humans is associated with genome-wide hypomethylation and some locus-specific hypermethylation (Jones et al., 2015), initial studies on cytosine methylation changes during leaf senescence and ageing had revealed contradictory evidence. In a variety of plant species and under different experimental conditions, cytosine methylation had been found either to increase or decrease (Dubrovina and Kiselev, 2016). Cytosine methylation is most important for transposon silencing, but also regulates gene expression, histone modifications and heterochromatin formation (Sequeira-Mendes et al., 2014; Lei et al., 2015). In plants, cytosine methylation is found in CG, CHG and CHH contexts, where H stands for nucleotides other than G (Stroud et al., 2013). In *Arabidopsis* methylation of some targets tended to be reduced during growth and ageing and this was correlated with reduced expression of the methyltransferase genes *MET1* (acting predominantly on CG) and *CMT3* (acting on CHG motifs), indicating less maintenance methylation (Ogneva et al., 2016). DNA demethylation can be either passive or active (Ikeda and Kinoshita, 2009). During DNA replication, the lack of maintenance can cause the passive loss of established methylation patterns, whereas active demethylation occurs enzymatically via base excision and repair by ROS1, DME, DML2 and DML3. ROS1 (REPRESSOR OF SILENCING 1) is particularly important for counteracting DNA methylation established by the RdDM pathway (Gong et al., 2002; Zhang et al., 2018). RNA-directed DNA methylation (RdDM) targets *de novo* methylation in all cytosine contexts and is triggered by the recruitment of DRM2 (reviewed by Matzke et al., 2015).

Detailed whole genome bisulfite sequencing (WGBS) on leaves during dark-induced leaf senescence in *Arabidopsis* revealed that the methylation landscape remains largely stable, with some hypomethylation occurring in CHH contexts (Trejo-Arellano et al., 2020). By contrast, Yuan et al. (2020) identified a genome wide reduction in methylation in all three cytosine contexts in age-related senescence, with the particular reduction of CG methylation in CG-rich sequences in close proximity to gene transcriptional start sites (TSS). Furthermore, the comparison with *dml3* knockout mutants identified a substantial correlation between cytosine methylation in the genes and their expression, suggesting an important role of the DML3-induced demethylation of gene regulatory regions during senescence (Yuan et al., 2020). However, despite clear evidence that methylation affects gene expression of some individual genes, a causal general relationship between global gene methylation patterns and transcript abundance is lacking (Zhang et al., 2018; Quadrana and Colot, 2016). Besides, natural age-related senescence and stress-induced premature senescence are known to exhibit major differences in, for example, their execution signalling and their transcriptome profiles (Buchanan-Wollaston et al., 2005). This may account for differences in methylation patterns in previous different experimental settings. Moreover, He and co-workers (2018) discovered a naturally occurring epiallele involved in the regulation of leaf senescence, additionally implying that cytosine methylation plays a role in senescence and in the adaptation of plants to local climates. Furthermore, progressive DNA demethylation occurs during tomato fruit ripening, with a profound effect on the fruit transcriptome (Lang et al., 2017). Finally, cytosine methylation regulates key genetic loci involved in flower induction: *FLOWERING LOCUS C* (*FLC;* Finnegan et al., 2005), *FLOWERING TIME* (*FT*; Zicola et al., 2019) and *FLOWERING WAGENINGEN* (*FWA;* Kinoshita et al., 2007). Although the key causal molecular link between flowering and the initiation of senescence has not yet been identified, cytosine methylation may play a role with its impact on flowering time and therefore affects the initiation of senescence. In addition to biotic stress (Dowen et al., 2012) and other abiotic stresses (Zhang et al., 2018), nutrient deficiencies elicit changes in the methylomes of *Arabidopsis* (Yong-Villalobos et al., 2015; Chen et al., 2018), maize (Mager and Ludewig, 2018) and rice (Kou et al., 2011). Thus, changes in the methylation landscape as a result of nutrient deficiency could (indirectly) have an impact on flowering time, senescence and nutrient remobilization.

In the present study, we analysed whether mutants that are defective in proper methylation exhibit altered development at the ultimate vegetative stages and during the generative growth phase. We employed the hypomethylated triple mutant *drm1 drm2 cmt3 (ddc)* and the hypermethylated *ros1* mutant in the genetic background of Col-0. We hypothesized that specific alterations of the methylation landscape are associated with the progression of generative growth and leaf senescence and performed WGBS at four time points, starting at the transition of the shoot apical meristem into reproductive growth. We further tested whether this developmental progress destabilizes the methylome and results in more variation. Our physiological, bioinformatic and biochemical analyses are consistent with moderately distinct cytosine methylation patterns at the different stages of leaf senescence in *Arabidopsis* that are not specifically related to senescence associated gene expression.

## Results

### Vegetative and reproductive growth, leaf senescence and nitrogen remobilization in *ddc* and *ros1*

The involvement of cytosine methylation in flowering time and leaf senescence was first investigated in the hypermethylated *ros1* (Gong et al., 2002) and the hypomethylated *drm1,drm2,cmt3* (*ddc*; Cao and Jacobsen, 2002) mutants. These mutants were grown in soil under long day conditions. Plants were scored for the transition to reproductive growth, for the first visible symptoms of senescence and for 50% senescence of the leaf rosette (Fig. 1A-C). Both mutants showed compromised rosette growth and smaller rosette diameter compared with Col-0, with the mutant *ros1* having the more severe phenotype (Suppl. Fig. S1). Aberrant methylation in these mutants tended to alter the flowering time and senescence progression, with the *ddc* mutant flowering slightly later and having delayed first symptoms of senescence. In these conditions, the *ros1* mutant tended to flower approximately 10 days earlier and exhibited earlier senescence compared with Col-0, although these differences in flowering time were only significant between the two mutants, but not between mutants and the wild type (Fig. 1A,B). Once the first symptoms of senescence had appeared, both mutants executed the senescence program faster than Col-0, with *ros1* being the more rapid (Fig. 1C). Whereas *ddc* had a similar total lifespan to that of Col-0, *ros1* had an overall shorter life time.

**Figure 1.**
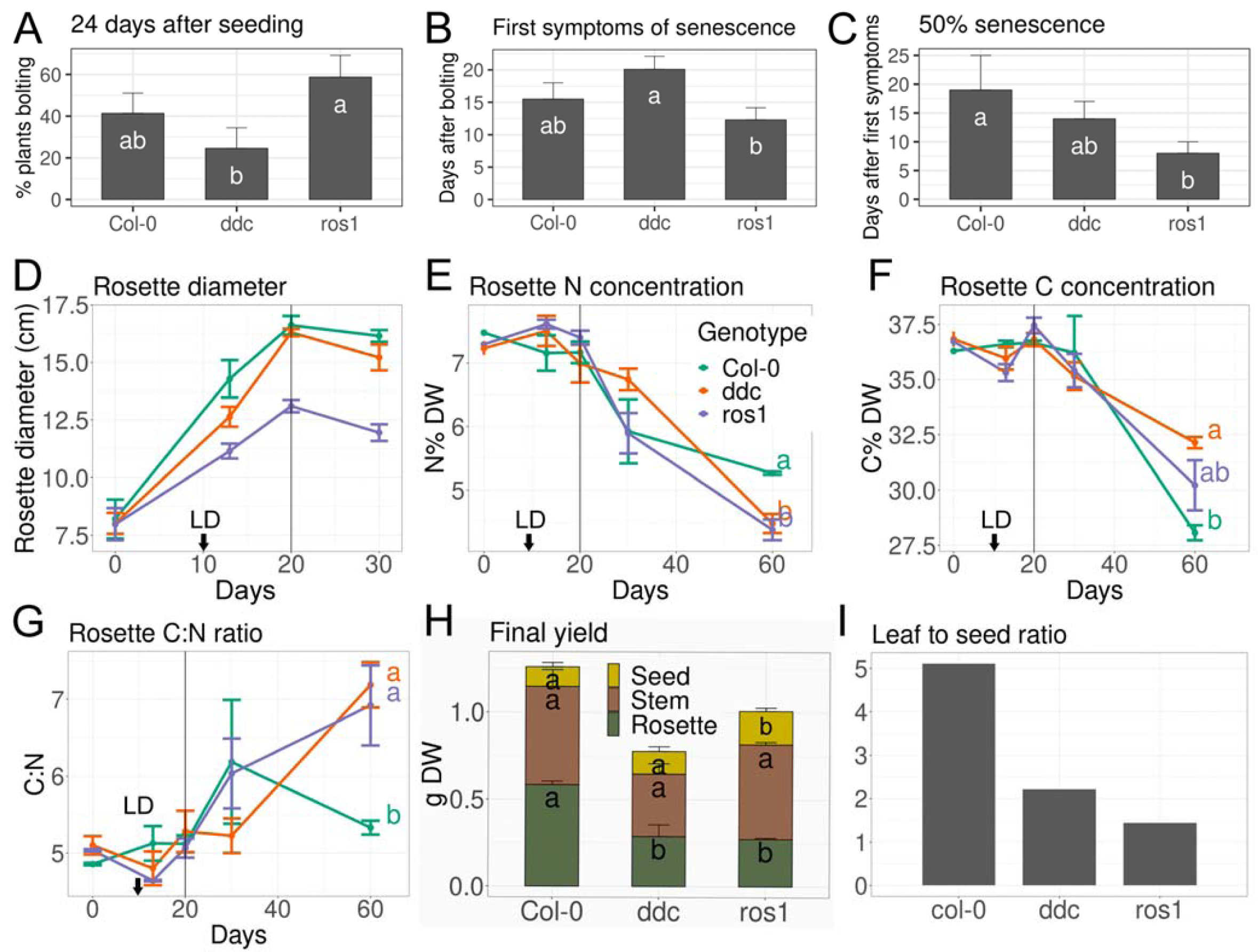
Flowering time and senescence of Col-0, *ddc* and *ros1*. (A) Percentage of plants that had undergone the transition of the shoot apical meristem (SAM) to flowering at 24 days after sowing in soil. (B) First symptoms of senescence of the whole rosette after appearance of the first bolt. (C) Days until 50% senescence (as visual count of the total rosette) was reached, since first symptoms were observed. (D-F) Time course for Col-0, *ddc* and *ros1* in hydroponics (sampling started at day 30 after seeding, this time point was set to 0 in the graph) with flowering induced via transition to long day (LD) regime at 10 days after first sampling (indicated by black arrow). All plants exhibited visual transition to reproductive growth in the shoot apical meristem at 20 days after first sampling (grey line). (D) Rosette fresh weight. (E) Nitrogen concentration in the rosette. (F) Carbon concentration in the rosette. (G) Carbon to nitrogen ratio of the rosette. (H) Rosette, stem and seed final dry matter yield. Statistical comparison was made separately for rosette, stem and seed biomass. Significant differences are indicated by different letters, p<0.05. (I) Leaf to seed ratio. Error bars indicate standard error; n=4; letters indicate significant differences estimated by LSD.test, following ANOVA at p< 0.05.

To examine possible physiological reasons and consequences of the aberrant senescence phenotype, carbon (C) and nitrogen remobilization during senescence were studied in these mutants in hydroponic culture. The plants were grown under short days, followed by a transition to long days (LD) to synchronize flower induction. The first samples were taken at 30 days after sowing. The three genotypes behaved as in the previous experiment, with Col-0 establishing the largest rosette biomass and *ros1* the least (Fig. 1D, Suppl. Fig. S1). In agreement with previous reports (Diaz et al., 2008), the N concentration in the rosette declined with proceeding senescence, but the initially delayed begin of senescence of *ddc* was correlated with a slightly later initiation of bulk nitrogen remobilization than in the other lines (Fig. 1E). Interestingly, *ros1* and *ddc* remobilized approximately 40 % more nitrogen than Col-0 out of the rosette until the end of the senescence process, when the leaves from the rosette had died (Fig. 1E). In contrast, the total carbon content of their leaves at the last harvest date was higher than that in Col-0 (Fig. 1F), corresponding to a higher overall leaf C:N ratio at the end of senescence (Fig. 1G). Final seed yield was higher in *ros1*, and *ddc* and *ros1* had less dry rosette biomass (Fig. 1H). This resulted in a lower overall leaf to seed ratio in senescent mutants, compared to Col-0 (Fig. 1I), suggesting that nutrient (nitrogen) remobilization and progression of senescence in *ddc* and *ros1* were affected, in agreement with the previous experiment.

### Flowering, leaf senescence and nitrogen remobilization with transient N shortage in *ddc* and *ros1*

As prolonged nitrogen deficiency was associated with massive overall decreases in cytosine methylation in maize roots (Mager and Ludewig, 2018) and maintained changes in the cytosine methylation patterns in the next generation were observed with differential N supply in rice (Kou et al., 2011), we tested the impact of temporary nitrogen withdrawal on the flowering time and senescence of the three genotypes grown in hydroponics. Nitrogen was reduced to 5% of the initial concentration (to 50 µM) for three weeks and was then resupplied. Flowering was induced by a change in the light regime (to long days, LD) at 10 days after N withdrawal. All three genotypes quickly reduced their growth rates, resulting in a reduced rosette biomass, because of low nitrogen (Fig. 2A). Col-0 produced significantly more rosette biomass under control conditions (Fig. 2A), whereas *ros1* produced the least biomass, especially when nitrogen was withdrawn. However, in contrast to Col-0, neither *ddc* nor *ros1* recovered the nitrogen concentration in the rosette directly following resupply (Fig. 2B). Nitrogen withdrawal delayed the flowering time in Col-0 but not in the two mutants and *ddc* showed a tendency to flower slightly earlier as a result of the treatment (Fig. 2C). Furthermore, *ros1* showed the highest seed yield, even after transient N deficiency (Fig. 2D). Again, *ros1* had the lowest N and highest C concentrations in the rosette at the final time point, corresponding to a higher C:N ratio (Fig. 2E).

**Figure 2.**
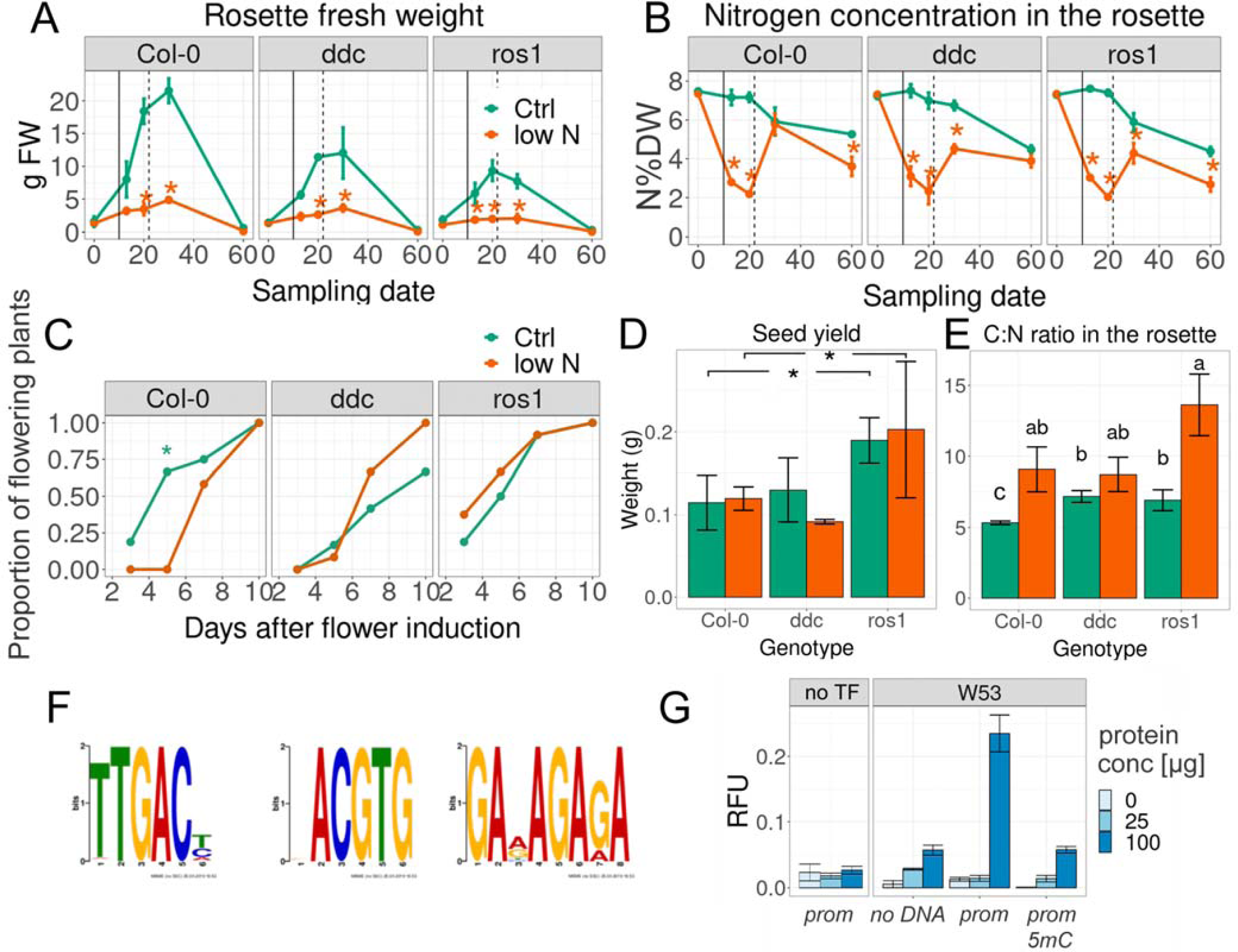
Temporary nitrogen withdrawal for 3 weeks in Col-0, *ddc* and *ros1,* and cytosine methylation targets in *ddc* and effects. Green: full nutrition, orange: withdrawal of N starting at day 0 (30 days old plants). On day 10, the light regime was changed; on day 21, the plants were resupplied with nitrogen. (A) Rosette fresh weights. (B) Nitrogen concentration in the rosette dry weight (N%DW). (C) Percentage of plants that visibly showed a bolt. The solid vertical line indicates induction of flowering via change in light regime and the dashed vertical line indicates nitrogen resupply. Significant differences are indicated by stars, p<0.05. (D) Seed biomass, (E) Carbon to nitrogen ratios in the rosette. Letters indicate significant differences as indicated by lsmeans test adjusted for TukeyHSD, following ANOVA at p< 0.05. Stars indicate significant effect of genotype only, without interactions between treatment and genotype (n=4; means ± SEM) (F) Top three most common transcription factor binding motifs found in DMRs between *ddc* and Col-0 intersecting with 2500bp promoters. E-values were: 1.5e-006, 56 and 2.5, respectively. (G) Cytosine methylation influence on the binding of WRKY53 via DPI-ELISA. Colour codes indicate the concentrations of protein (in µg per 60 µl reaction), “*prom”* (concentration: 0.33 pmol * µl^-1^) non-methylated, artificial promoter and “*prom 5mC”* methylated, artificial promoter (n=2; means ± SEM).

### Cytosine methylation in WRKY binding sites is functionally important and affected in *ddc*

We considered the possibility that the aberrant remobilization and senescence phenotype in *ddc* was caused by an interference of cytosine methylation in transcription factor binding sites near genes. To test this hypothesis, we analysed the methylome datasets of *ddc* and Col-0 provided by Stroud et al. (2013). *ddc* had 18% less CG methylation and 89% and 56% reduced methylation in the CHG context (in which H means any base out of A,T,C) and in the CHH context (Suppl. Fig. S2A), respectively. This led to 228 differentially methylated regions (DMRs) of which 186 intersected with annotated genomic elements, the majority being found in transposable elements (TEs) or -2500bp from the transcription start site (TTS) of gene open reading frames (ORFs) (Suppl. Fig. S2B). 178 motifs were recognized within DMRs that intersected with 2500bp promoter regions. The most prevalent group was *TTGAC(T/C/A*), corresponding to the conserved W-box element recognized by WRKY transcription factors (TFs, Fig. 2F). The second most differentially targeted motif was part of a core motif recognized by bZIP transcription factors (*ACGTG*), opening the possibility that the loss of methylation in CHG and CHH contexts affected the WRKY and/or bZIP transcription factor network.

Using artificial binding targets, we verified that methylation massively inhibited *in vitro* DNA binding of key transcription factors of senescence, namely WRKY53 (Fig. 2G), WRKY18 and WRKY25 (Suppl. Fig. S3), by biochemical binding assays (Brand et al., 2010). Whereas WRKY18 bound the most strongly to the artificial 3x W-box target when non-methylated (Suppl. Fig. S3A,C), the interaction of all three proteins with the artificial promoter sequence was greatly impaired by (complete) cytosine methylation of the artificial target (Fig. 2G, Suppl. Fig. S3).

### Relationship between reproductive stage, rosette size, leaf coloration, chlorophyll and leaf physiology in Col-0

In order to analyse genome methylation changes within individual leaves along the senescence process, we chose four physiologically relevant time points in the reproductive development of the plant: Bolting (Blt), Flowering (Flwr), Seed Development (SD) and Seed Maturation (SM) (Suppl. Fig. S4). Leaves were marked with coloured threads shortly after their appearance so that, even at later stages, the leaves were distinguishable and could be numbered according to their age (with number 1 being the oldest and first emerging true leaf). Leaves numbered 5 to 9 exhibited sequential senescence, with leaf number 5 showing the earliest symptoms attributable to chlorophyll degradation, namely as early as the initiation of flowering (Fig. 3A; Suppl. Fig. S5). All leaves enlarged and gained weight until Flwr, the second time point (Fig. 3B). An automated colorimetric assay (ACA; Bresson et al., 2018) trained on leaf number 5 precisely predicted chlorophyll concentrations (adjusted R^2^ = 0.89) in leaves numbered 6 to 9. The colorimetric and visible loss of green coloration and its variation (Fig. 3A, Suppl. Fig. S5) were largest at the third time point (SD), as indicated by principle component analysis (PCA, Fig. 3C, Suppl. Fig. S5). By normalizing leaf weight to a scale between -1 and 1, different leaves of distinct plant individuals were successfully assigned directly to certain stages, despite their slightly disparate size and time after seeding due to biological variation (Suppl. Fig. S5). A peak of hydrogen peroxide was observed at the flowering stage, when leaves numbered 6 and 9 approached their maximum size and weight (Fig. 3D; Suppl. Fig. S5). Glucose increased as early as Flwr, when the first reduction in chlorophyll occurred (Fig. 3E; Suppl. Fig. S5), followed by an increase of fructose at SD, when glucose concentrations peaked (Fig. 3E).

**Figure 3.**
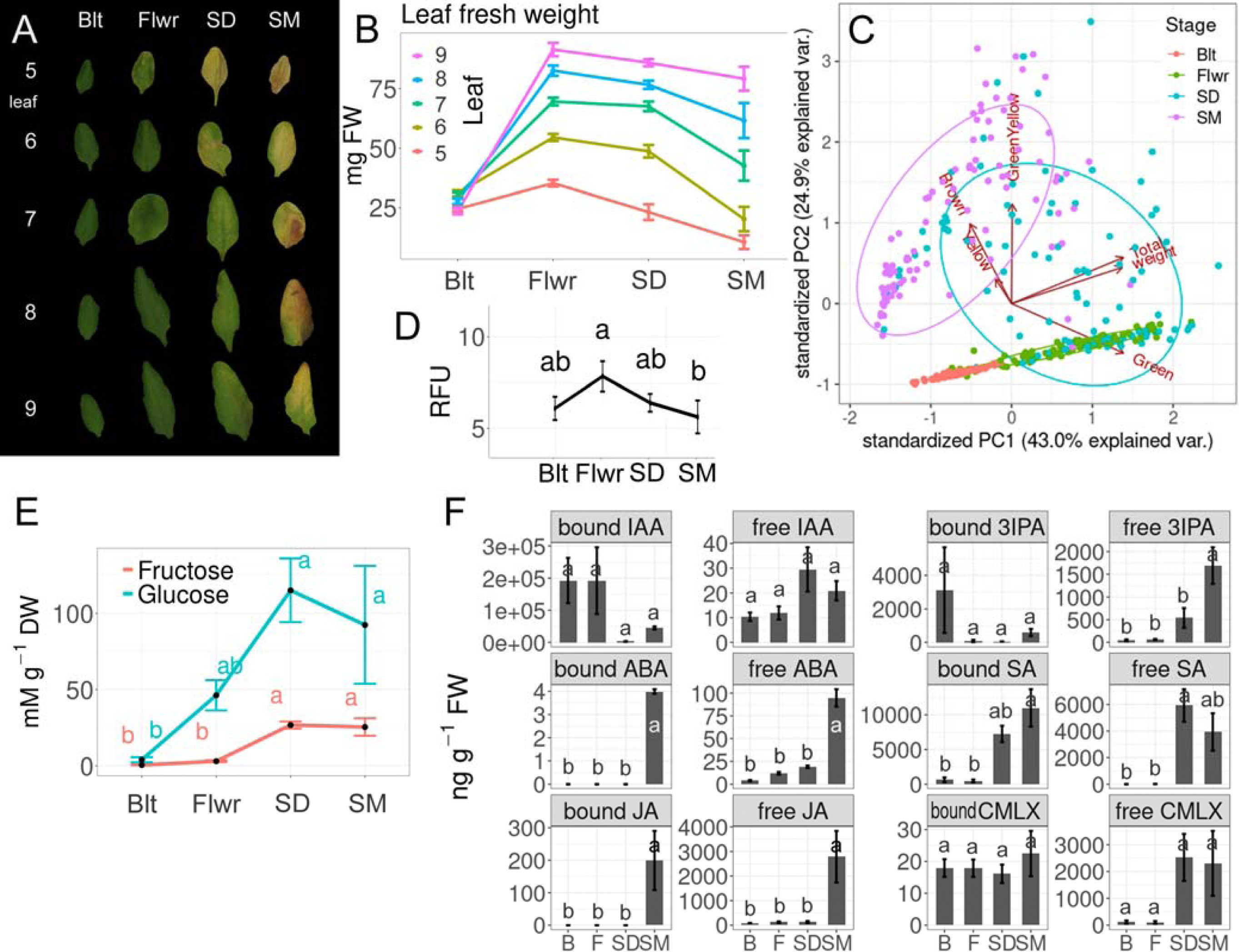
Progression of senescence in leaves numbered 5 to 9. (A) Fresh weights measured on per leaf basis (n = 20; means ± SEM). (B) PCA on pixel counts for green, green-yellow, yellow and brown coloration, total pixel count and leaf fresh weight distinguishing the four developmental stages: Bolting (Blt/B), Flowering (Flwr/F), Seed development (SD) and Seed maturation (SM). (C) Representative leaves at the four stages. (D) Hydrogen peroxide in leaf nine. RFU = relative fluorescent units. (n = 20; means ± SEM; LSD.test following ANOVA) (E) Glucose and fructose concentrations in leaf number 6 (n = 4; means ± SEM; LSD.test following ANOVA). (F) Concentration of hormones, precursors and defence-related compounds: auxin (IAA, 3-IPA), abscisic acid (ABA), salicylic acid (SA), jasmonic acid (JA); and camalexin in soluble (free) and hydrolysable (bound) form (n=3; means ± SEM).

Leaf number 8 was then analysed for various phytohormones, namely the auxin indole acetic acid (IAA), its precursor indole-3-pyruvate (3-IPA), abscisic acid (ABA), salicylic acid (SA), jasmonic acid (JA) and defence-related anthocyanins and camalexin. Whereas the phytohormones gibberellic acid, auxin and cytokinin are considered to inhibit senescence, abscisic acid (ABA), jasmonic acid (JA) and salicylic acid (SA) are inducers of senescence (Jibran et al., 2013; Schippers et al., 2007). Hydrolysable (bound) IAA tended to decrease between Flwr and SD, while free IAA tended to increase at this time point (Fig. 3F) and its precursor 3-IPA was only elevated at SM (Suppl. Fig. S6). Free SA also increased between Flwr and SD (Fig. 3F). This occurred after the intracellular hydrogen peroxide had reached its peak (Fig. 3D) and was paralleled by an increased amount of the anthocyanins calabricoside A, cyanidin and kaempferol (Fig. 3G), as well as the defence-related phytoalexin camalexin in its free form (Suppl. Fig. S6). This indicates that the progression of senescence was associated by the activation of defence mechanisms of the plant between Flwr and SD, before JA and ABA increased. These two phytohormones increased later at SM, both in their bound and active free forms (Fig. 3F).

### Time course of cytosine methylation changes in leaf 7 during ageing and senescence

Because of the phenotypes of the two methylation mutants (Fig. 1, 2) and the inconsistent methylome changes reported previously to be associated with leaf senescence (Trejo-Arellano et al., 2020; Yuan et al., 2020) we decided to analyse whole genome cytosine methylation at the four stages (Blt, Flwr, SD and SM) in leaf number 7. High conversion efficiency and the sequencing measures indicated the high overall quality of our data (Suppl. Table S2). Use of triplicates ensured sufficient statistical power for the subsequent analyses and allowed an estimate to be made of the rate of stochastic variation in the progression of leaf decay. A total of 45,5047 differentially methylated loci (DMLs, 25% change) were identified in the chromosomal DNA, with approximately 78% being unique between the six pairwise comparisons of stages. Most DMLs were found in the CHH context (169015), followed closely by the CG context (163909), with the least DMLs in the CHG context (122123).

To estimate the random noise and stochastic variation of cytosine methylation and the ability of the statistical package to pick stage-defining DMLs, we performed clustering and PCA on the methylome data with 50 bp windows with weighted means (Fig. 4). We first questioned whether each methylome dataset was strictly associated with the chosen individual developmental stage, i.e. whether triplicate datasets from each developmental stage clustered together, but separately from the others. This analysis revealed two outliers of the complete methylome datasets at Blt and SM, which had less technical sequencing quality than the other samples (Suppl. Table S2). These were grouped together in the cluster analysis and separated the samples in PCA component 1 (Fig. 4A,C). DMLs of the four stages, however, were well-defined on both graphs, allowing us to select against random noise to pick stage-defining DMLs (Fig. 4B,D). DMLs appeared throughout the chromosome but the densest DML regions with higher average differential methylation were found around the centromere. In this region the highest concentration of TEs was found (Suppl. Fig. S7). There was progressively more hypo-than hypermethylation between Flwr and SD, but this reversed with the maturation of the seeds (SM) (Fig. 4E). The total average cytosine methylation changes were small overall, but methylation dropped initially in all contexts examined and reached a minimum at SD (Fig. 4E). This drop was strongest in the CG context. The last stage of chlorophyll degradation and the beginning of desiccation during SM corresponded to a period of moderate *de novo* methylation that recovered to total pre-flowering methylation levels. Interestingly, however, the CG context did not recover completely, whereas cytosines in the CHG and CHH contexts in TEs and 2500bp promoters reached levels higher than those beforehand (Fig. 4E). The largest reductions in methylation were observed in the CG context in TEs. Inhibition of the maintenance methylation during cell division and active RdDM might have been responsible for this finding and are in agreement with such patterns.

**Figure 4.**
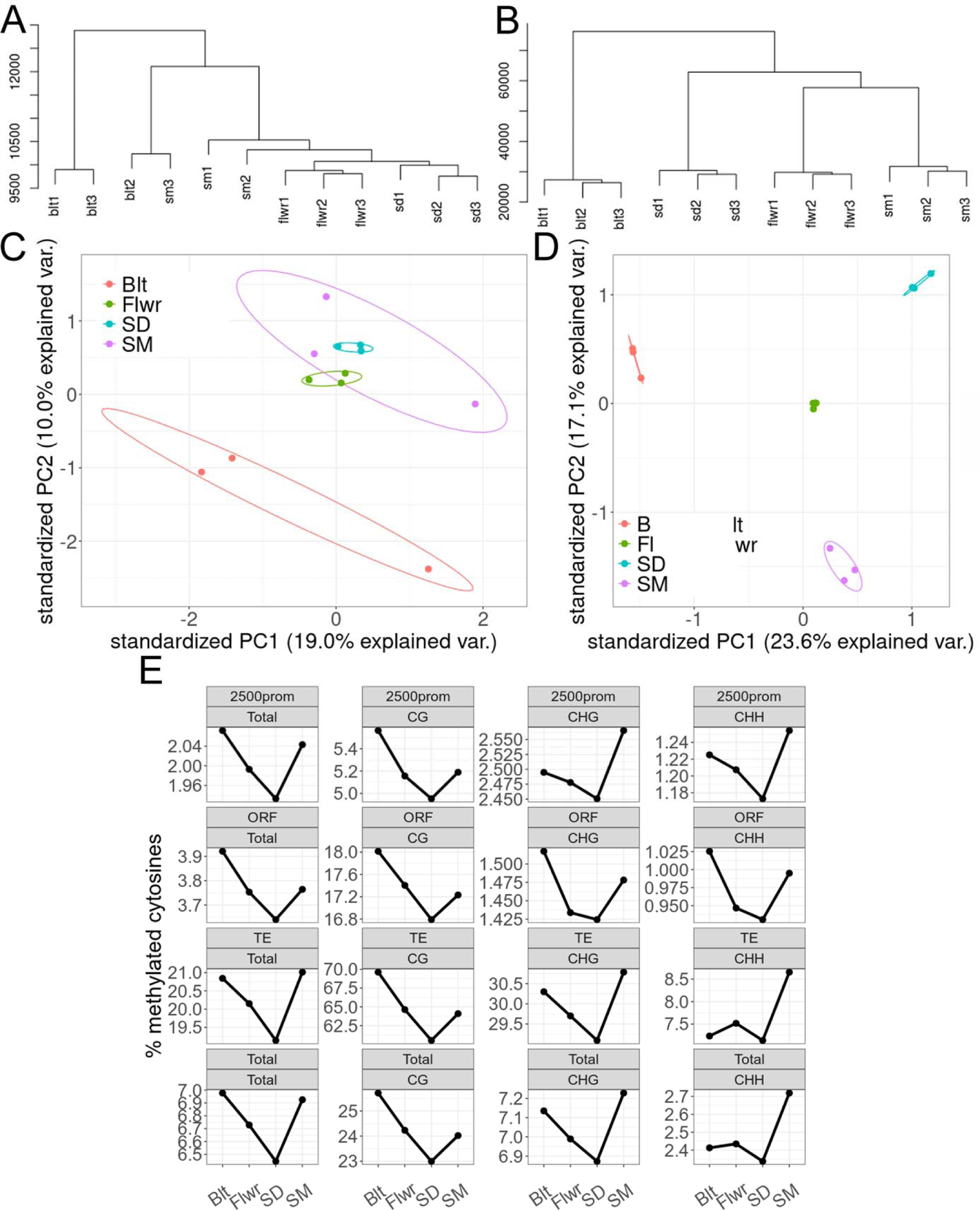
Cytosine methylation in leaf number 7. (A-D) Reproducibility of the WGBS experiment. (A) Cluster dendrogram of the mean methylation calculated at 50bp bins. (B) Cluster dendrogram of all cytosines categorized as DMLs in at least one pairwise comparison. (C) PCA analysis of the mean methylation calculated at 50bp bins. (D) PCA analysis of all cytosines categorized as DMLs in at least one pairwise comparison. (E) Methylation of cytosines in 2500bp promoter regions, open reading frames (ORF), transposon elements (TE), and total methylation of the whole genome (total) during four developmental stages, analysed independently of context and separately in all three cytosine contexts (CG, CHG and CHH).

Differential methylation occurred mostly in TEs, followed by ORFs, and was found to be lowest in 2500bp promoters (Fig. 5A). Interestingly, a large proportion of TEs containing DMLs intersected with ORFs and their 2500bp promoters (Fig. 5A). About half of the loci that were hypomethylated between Blt and Flwr were in the CG context and one quarter was in CHG and CHH contexts, respectively. About two thirds of all hypomethylated loci were in the CG context (Fig. 5B). In contrast, hypermethylation predominantly occurred in CHG and CHH contexts throughout the observed stages (Fig. 5C). Stage-by-stage comparisons revealed gradual changes between stages and that hypomethylation dominated between Blt, Flwr and SD, whereas hypermethylation dominated between SD and SM, with about three quarters of DMLs being hypermethylated (Fig. 5D).

**Figure 5.**
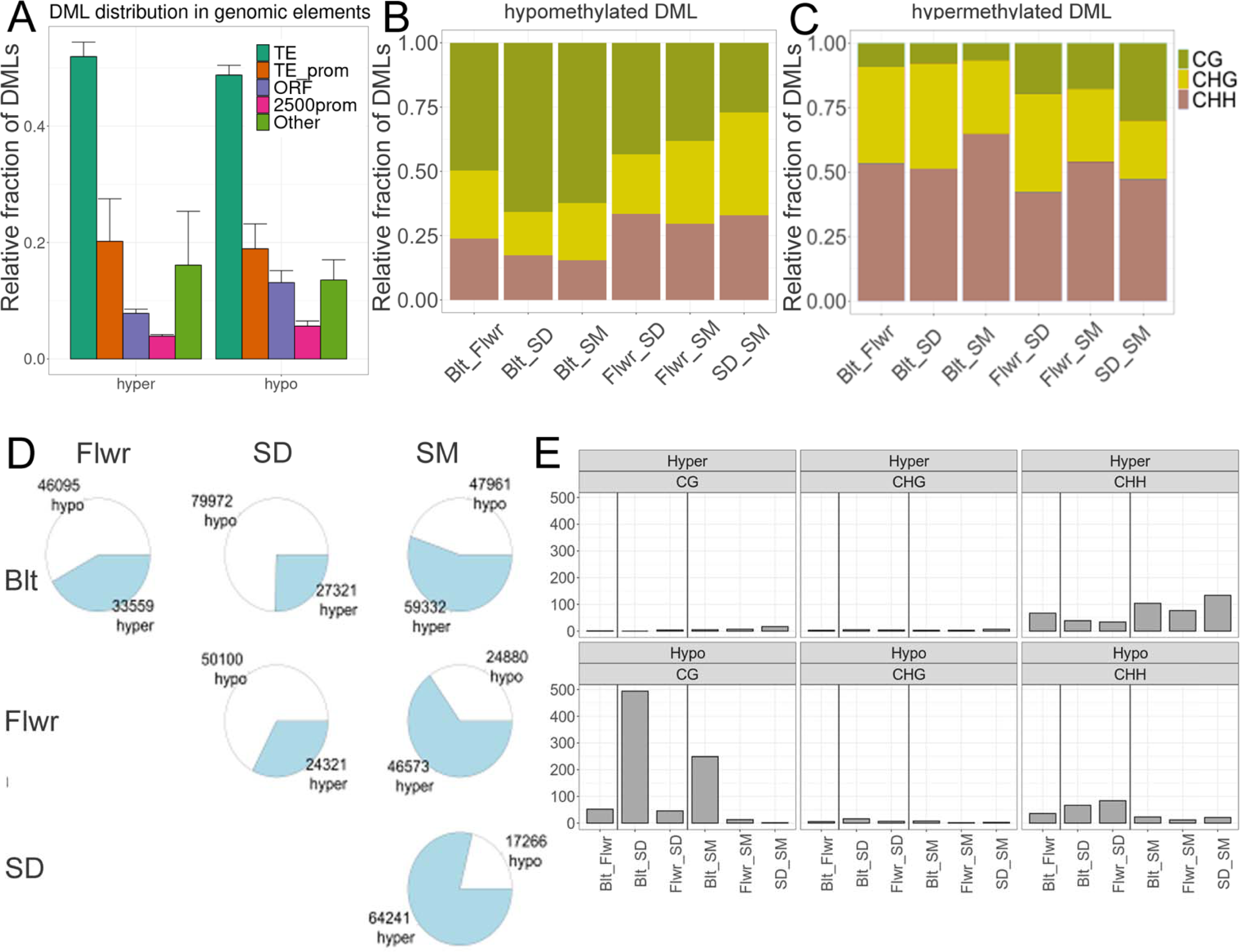
Relative distribution of differentially methylated loci (DML) in genomic contexts, hypo- and hypermethylated loci and differentially methylated regions (DMRs). (A) Proportion of DMLs in TEs outside of promoters (dark green), TEs within 2500bp promoters (orange), ORFs (violet), 2500bp promoters that lack TEs (magenta) and intergenic regions (light green). Error bars indicate SEM. (B) Proportion of hypomethylated and (C) hypermethylated DMLs in CG (green), CHG (orange) and CHH (violet) contexts. (D) Pairwise comparison of hypo- (white) and hypermethylated (light blue) loci in all pairwise comparisons. (E) DMRs in the six pairwise comparisons separated by cytosine context.

### Distribution and characteristics of differentially methylated regions (DMRs)

We found a relatively small number of differentially methylated regions (DMRs, definition see methods) associated with the different leaf stages, which precisely followed the patterns of DMLs (Fig. 4E, Fig. 5E). Most DMRs (494) were observed in the CG context between Blt and SD, followed by Blt and SM (251). We noted that these numbers were not the sum of DMRs found between individual time point comparisons (each <50), meaning that DMRs developed gradually with time as a superposition of contexts (Fig. 5E). DMRs in CHH (total number: 683) were primarily hypomethylated until SD, whereas hypermethylated CHH DMRs were present throughout the experiment. DMRs in the CHG context were negligible throughout the process (Fig. 5E).

In order to recognize genes that were potentially influenced by DMRs with regard to their gene expression, we selected all ORFs and 2500bp upstream regions containing DMRs. 132 genes had at least one DMR intersecting with their ORF (Suppl. Table S3) but no significant enrichment was found when we probed the geneontology.org database or senescence-associated genes (Breeze et al., 2011). Some notable groups included 10 genes involved in lipid metabolic processes, 12 genes involved in the hormonal response, 25 genes involved in stress responses, 16 genes involved in phosphorus metabolism and 13 genes involved in protein phosphorylation. Interestingly, three WRKY transcription factor genes were found with DMRs located in their ORF, namely *WRKY18, WRKY26* and *WRKY50*. Moreover, *CYTOKININ OXIGENASE/DEHYDROGENASE 1 (CKX1)*, which codes for the protein involved in catalysing the degradation of cytokinins, was in this group. A further 581 genes were found with at least one DMR intersecting with the region 2500bp upstream of their TSS (Suppl. Table S4). Again, no significant enrichment was found when we probed databases, such as geneontology.org. Of these genes, 25 were involved in reproduction, 11 in the posttranscriptional regulation of gene expression, 42 in cellular component organization or biogenesis, 21 in peptide metabolic processes and 10 in methylation. Genes worth mentioning with DMRs in the promoter were *ROS1*, *SUVH1,* which is involved in methylation recognition and the phytochrome photoreceptor gene *PHYC*. The glutamine synthetase gene *GLN2*, three WRKY transcription factor genes (*WRKY41, WRKY48* and *WRKY64*), two bZIP transcription factor genes (*bZIP23, bZIP60*), five NAC transcription factor genes (*NAC001, NAC003, NAC019, NAC071, NAC100*) and one MYB gene (*MYB62*) also had DMRs in their promoters. None of these genes is, however, considered as a central regulator of senescence, of flowering or of pathogen defence, with the exception of *NAC019*, that is potentially involved in senescence.

### Cytosine methylation and gene expression

Although our previous analyses did not support any strong direct links between differential cytosine methylation and senescence-associated gene expression, we performed *qRT-PCR* on the same leaf samples as those used for WGBS to ascertain the temporal relationships between cytosine methylation and gene expression for at least a few genes. The promoter region of *ROS1 (AT2G36490)* appeared to be the most interesting because it was hypomethylated in the early stages of flowering, but methylation partially recovered at SM (Fig. 6A). The location of this transient DMR corresponded to a genomic position that was previously recognized as a DNA <u>me</u>thylation <u>m</u>onitoring <u>s</u>ystem involved in gene methylation regulation via *ROS1* (MEMS; Lei et al., 2015; Zhang et al., 2018). Less methylation in all three cytosine contexts was observed in the *ROS1* MEMS region (Fig. 6A) between Blt and Flwr, which coincided with reduced *ROS1* expression, as expected from the nature of this region (Lei et al., 2015).

**Figure 6.**
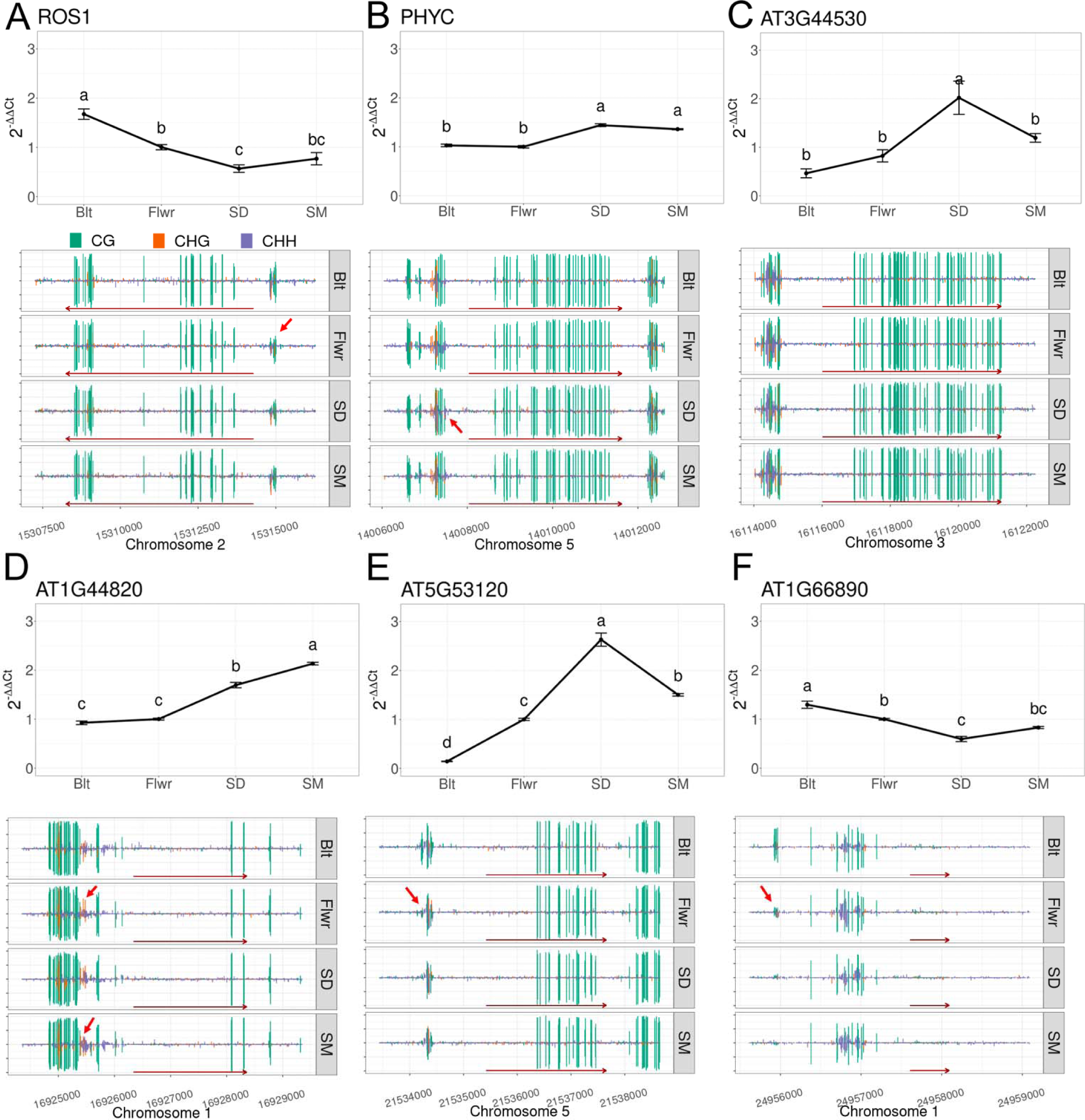
Patterns of gene expression (qRT-PCR) and cytosine methylation (CG, green; CHG, orange; CHH, violet) in representative genes. (A) *ROS1,* (B) *PHYC*, (C) *AT3G44530* (no DMR in this gene and promoter), (D) *AT1G44820,* (E) *AT5G53120* and (F) *AT1G66890* for time points Blt, Flwr, SD and SM. Dark red arrows along the x axis of the gene methylation graphs indicate the gene span and direction. The small red arrows point at differentially methylated regions (DMRs). For gene expression analysis, two pooled biological samples with 3 technical replications and at least 2 repetitions were used. Error bars indicate SEM.

Another DMR found upstream in close proximity to the phytochrome *PHYC* gene spanned 16 nucleotides and occurred only in the CHH context. Less methylation was observed between Flwr and SD (Fig. 6B), coinciding with a small increase in *PHYC* expression. However, several senescence up-regulated genes, such as *AT3G44530* (encodes a WD-containing component of a putative HISTONE chaperone complex), *AT1G44820* (codes for a putative aminoacylase), *AT5G53120* (encodes a putative spermine synthase) and *AT1G66890* (unknown function) that were associated with massive cytosine methylation losses in their promoters in previous analyses during senescence (Yuan et al., 2020), had more diverse methylation patterns in our study (Fig. 6C-F). For *AT3G44530,* a gene up-regulated at early stages, no changes in cytosine methylation were detected in our dataset. Methylation in the *AT1G44820* promoter increased transiently, while up-regulation of gene expression increased at early stages. The methylation in the *AT5G53120* promoter increased at early stages, whereas gene expression transiently increased until SD and then decreased. A slight reduction in *AT1G66890* expression levels until SD was associated with the loss of a cytosine methylation DMR in its promoter before flowering (Fig. 6F). Any simple, causal temporal relationship between methylation in gene promoters and SAG expression regulation appears unlikely for this small set of genes.

### Cytosine methylation in bZIP and WRKY transcription factor binding sites

We therefore considered the possibility that DNA/gene transcription factor interaction networks as a whole might be affected by differential cytosine methylation at many unrelated loci. Cumulative effects within gene expression networks might then only be expected when several transcription factor binding sites are targeted by differential methylation in certain motifs, as in the case of the *ddc* mutant (Fig. 2E). Indeed, we found 2300 DMLs inside bZIP binding sites, of which 71 coincided with DMRs, and 3892 DMLs in W-boxes, of which only 25 matched with DMRs. Differentially methylated transcription factor binding sites were, however, only a minor fraction of genome-wide sites; they represented only 3 % and 1.7 % (of a total 73098 bZIP and 235305 WRKY sites), respectively.

We noted that the methylation of *bona fide* bZIP binding sites that contained A-, C- or G-box palindromes with a CG motif was depleted in all genomic regions, in promoters, in ORFs and TEs. The largest differences were observed in promoter regions 2500bp upstream of TSS (Fig. 7A). Despite the large number of differentially methylated bZIP sites, average temporal methylation changes at these sites were small and declined until the SD stage. In contrast, the conserved six-nucleotide sequence that is targeted by WRKY transcription factors (known as the W-box, *TTGACT/C*), had a different methylation status in TEs compared with the overall methylation status of cytosines in the same context in other regions (Fig. 7B), but again, the average temporal variation was small. The CHH context of the W-box (*TTGA**C**T**H*** and *TTGAC**CHH***) had a significantly higher methylation level in promoters, ORFs and TEs, but the overall occurrence of CHH methylation in W-boxes in 2500bp promoters was the lowest and its significance to be different from all regions declined during senescence (Fig. 7B). ORFs and TEs maintained higher methylation throughout the time course. The CHG context appeared less often in a W box (*TTGA**CTG*** and *TTGA**CCG***) and had higher methylation only in TEs. Even though the two groups of transcription factor binding sites exhibited opposite behaviour in terms of cytosine methylation, their presence in or close to ORFs was consistently associated with lower methylation levels, compared with other genomic regions.

**Figure 7.**
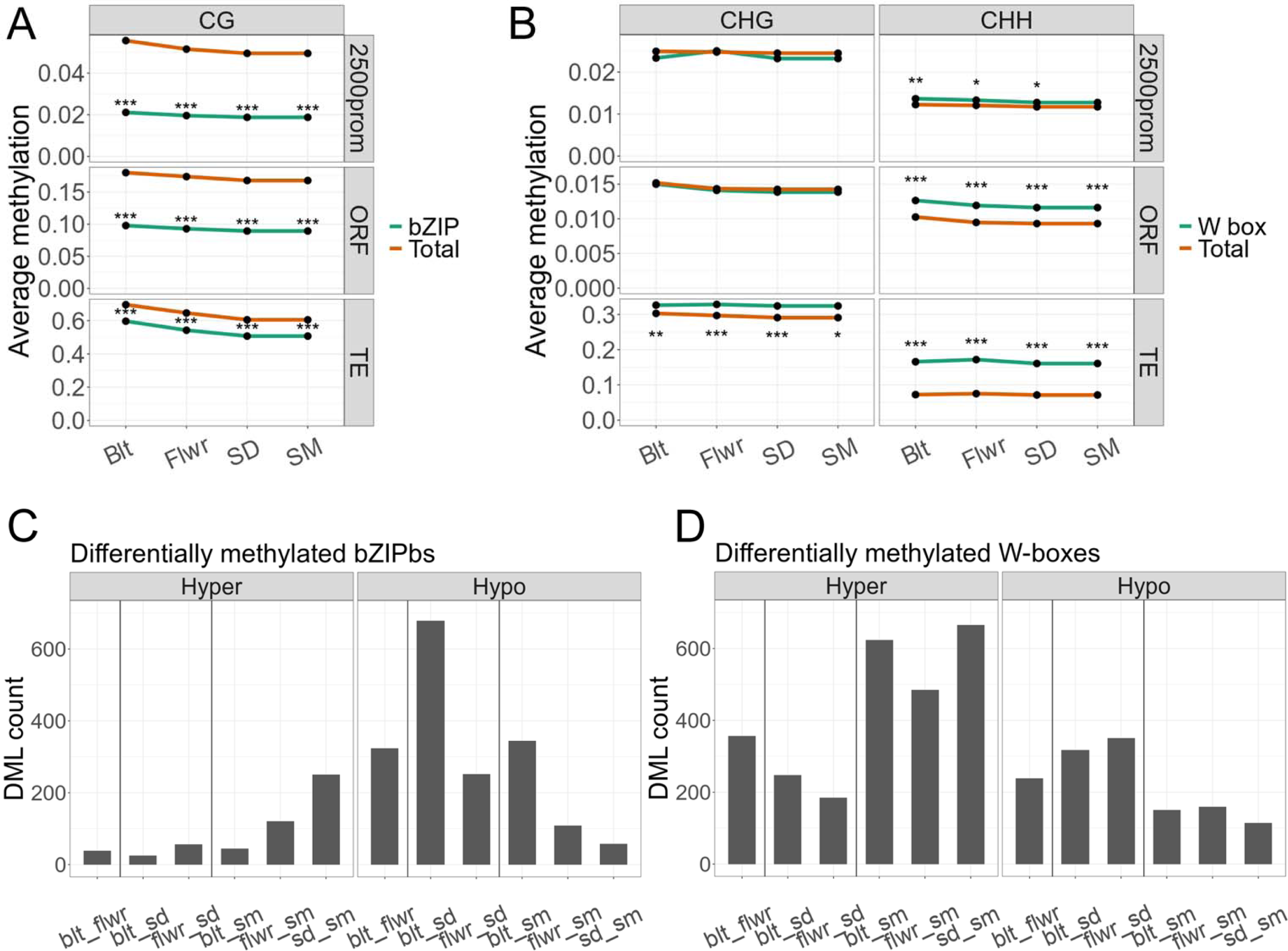
Average methylation levels of bZIP and WRKY transcription factor binding sites (green) compared to overall genome methylation (orange) in regions 2500bp upstream of TSS, ORFs and TEs in their respective cytosine context. (A) Average methylation of A-, C- and G-boxes. (B) Average methylation of W-boxes. Bootstrap analysis was performed. Significance levels: * <0.01; ** <0.001; *** <0.0001. (C) Number of A-, C-, G- or (D) W-boxes intersecting with DMLs.

In total, 358 genes were found with differentially methylated A-, C- or G-boxes (Suppl. Table S5). Significant over-representation was found in only one group, comprising 7 genes related to the regulation of gamete fertilization, all of them coding for ECA1 gametogenesis-related family proteins. Interestingly, a differentially methylated A-box was located in the vicinity of the *PHYC* gene (Fig. 6B), together with a differentially methylated G-box near *NAC019*. 517 genes were found with a differentially methylated W-box in their proximity (Suppl. Table S6), but no significant over-representation (and no relationship with senescence-associated genes) was found in this group. Notable genes were *WRKY74* with three differentially methylated W-boxes, and *WRKY41* with two differentially methylated W-boxes. *WRKY64, NAC100, NAC003* and *RDR1* contained one differentially methylated W-box. Even though none of these genes are recognized as major regulators of senescence, this list agrees with the potential involvement of differential methylation in transcription factor binding networks.

## Discussion

The involvement of epigenetic regulatory mechanisms in plant longevity, leaf ageing and senescence has been previously established (Ay et al., 2009). Here, we initially tested whether the aberrant maintenance of cytosine methylation in the *ddc* and *ros1* mutants resulted in senescence- and remobilization-specific phenotypes. Clearly, these mutants have pleiotropic developmental defects (with flowering initially reported to be unaffected by non-CG methylation; Chan et al., 2006) and we cannot exclude that methylation changes are secondary. However, in our conditions the hypomethylated *ddc* mutant flowered later than *ros1* and had delayed first symptoms of senescence compared to hypermethylated *ros1* (Fig. 1). The execution of the senescence program following the appearance of the first symptoms tended to be faster in both mutant lines compared to the wild type and resulted in altered leaf to seed ratio and nitrogen remobilization from the rosette and an overall higher C:N ratio at the end of the process. Diaz et al. (2008) found similar N partitioning between sink and source tissues, but this did not correlate with early or late senescence phenotypes. Clearly, methylation mutants have pleiotropic phenotypes and cytosine methylation might indirectly alter the C:N ratio or hormonal regulation. Indeed, increased IAA concentrations were observed in the leaves of *ddc* seedlings, consistent with differences in the expression of genes involved in auxin synthesis, transport and signalling (Forgione et al., 2019).

Furthermore, proper maintenance of cytosine methylation was required during N limitation. While transient nitrogen withdrawal caused significant reductions in rosette biomass in all three genotypes, the two mutants did not completely recover transiently after N resupply and showed lower terminal N concentrations in the rosette compared with Col-0 (Fig. 2). In accordance with the findings of Lin and Tsay (2017), N limitation delayed flowering in Col-0, but this was not seen in the methylation mutants (Fig. 2). Furthermore, both mutants apparently more efficiently remobilized N to seeds, which might be interesting from an agronomical point of view, if confirmed in crops. *ros1* produced more seed biomass, regardless of the treatment. Lower leaf to seed ratios, together with higher N remobilization and higher C:N ratios in the rosette under control conditions agree with faster senescence progression and differential N remobilization in *ddc* and *ros1* (Fig. 1, 2). Interestingly, Kuhlman et al., (2020) have suggested an involvement of cytosine methylation in the regulation of specific loci related to shoot growth under N deficiency in *Arabidopsis*. Cytosine methylation changes associated with leaf senescence, however, did not specifically target N-related genes or their promoters in our experiments (Suppl. Tables S3, S4, S5, S6). Nevertheless, several *NAC* and *WRKY* transcription factor genes including one involved in senescence, namely *NAC019* (Hickman et al., 2013; Takasaki et al., 2015), were differentially methylated, implying that these might gradually orchestrate changes in transcription factor networks.

We chose four physiologically relevant stages and five distinct leaves per plant for various analyses to relate these to ongoing leaf senescence. Colour, size and weight correlated with chlorophyll concentrations (Fig. 3, Suppl. Fig. S5). In close agreement with previous reports, net loss in chlorophyll concentration (due to repression of its biosynthesis starting around the induction of flowering at 21 days after seeding; Breeze et al., 2011) occurred as soon as the transition of the shoot apical meristem to reproductive growth occurred. This was accompanied by increases in soluble sugar content (Wingler et al., 2006), elevated hydrogen peroxide levels and a moderate overall reduction in cytosine methylation (Fig. 4). Comparing leaf material at certain stages (instead of sampling at fixed days after seeding) decreased the variance within sampling groups, because the senescence time course of plant individuals is subject to substantial variation even in a single genotype. Differences in flower induction of the two methylation mutants (Fig. 1,2) are likely explained by distinct cytosine methylation of crucial genes and their promoters (Finnegan et al., 2005; Zicola et al., 2019; Kinoshita et al., 2007). Hormonal changes during leaf senescence appeared later in the process (Fig. 3, Suppl. Fig S6) and identified a strong activation of defence reactions. Increases in SA and phytoalexins at SD preceded elevation of JA and ABA. It is interesting to note that widespread DNA methylation changes are associated with biotic stress responses (Dowen et al., 2012) and that such stress responses were prominent at the terminal stages, at which *de novo* methylation occurred (Fig. 4).

We found moderate methylome changes and an interesting discrepancy between differential methylation in CG, CHG and CHH contexts; more than 50% of all hypomethylated loci appeared in the CG context (Fig. 4, 5). This trend shifted towards CHG and CHH between SD and SM. On the other hand, hypermethylation occurred predominantly in CHG and CHH contexts, with a slight shift towards CG during SM. Previous studies suggested that the two main methyltransferase genes involved in CG methylation maintenance, namely *MET1* and *CMT3*, are down-regulated after flower induction in *Arabidopsis* (Breeze et al., 2011; Ogneva et al., 2016). Leaf growth ceased around Flwr, so the hypomethylation in CG is at least partially attributed to the reduced maintenance of methylation following genome duplication, as leaf cells in *Arabidopsis* divide almost until the final leaf size is reached, with endoreduplication occurring from the time that cell division rates decline, until the end of cell expansion. The relatively low amount of differentially methylated regions is clearly also due to the fact that even at the defined sampling times, leaf cells within the same leaf are visibly in different stages. Clearly, senescence progression from the tip is not entirely uniform in every leaf (Fig. 3A,C) and it might be necessary to resolve stage-dependent methylomes at the cellular level. Our 25% criterium for DMLs and DMRs was chosen to account for discrepancies between cell stages in the same leaf; differential methylation at 25% likely reflects a 100% change in the methylation of a quarter of all leaf cells (as methylation is either present or not). It may be possible (and necessary) in the future to separate different cellular leaf senescence stages by cell sorting of enzymatically isolated leaf cell protoplasts. Still, hypomethylation was clearly not concentrated in specific genomic regions, but rather appeared throughout the genome. The partial recovery of CG methylation after the cessation of leaf expansion, between SD and SM, might be the result of residual activity of maintenance methyltransferases. However, the large number of hypomethylated loci when Blt is compared with SM indicates that this increase is caused by *de novo* methylation, rather than the recovery of lost methyl-cytosines. Accordingly, the main methyltransferase genes involved in CHG and CHH methylation, namely *DRM1* and *DRM2*, showed (mostly) stable expression during senescence (Breeze et al., 2011; Ogneva et al., 2016). This suggests a functional role of RdDM and *de novo* methylation in leaf senescence progression as its loss affects senescence progression and agrees with our finding that the triple mutant *ddc* has an aberrant senescence and remobilization phenotype (Fig. 1, 2).

Interestingly, dark-induced leaf senescence, a process more similar to the ultimate senescence steps between SD and SM, was associated with moderate hypomethylation in the CHH context (Trejo-Arellano et al., 2020), demonstrating differences between the faster, stress-induced senescence and the slowly progressing age-related type of senescence studied here. Similar to our study, the methylation changes in the CHH context in dark-induced senescence did not correlate with senescence-associated gene expression (but were associated with the up-regulation of young transposons; Trejo-Arellano et al., 2020). In agreement with that study, the ultimate stages in our experimental setup also consistently targeted predominantly the CHH context, which is again in agreement with ongoing RdDM. In contrast, Yuan et al. (2020) have described similar patterns of CG demethylation during initial phases of natural senescence in leaves, as observed here (Fig. 3), but did not observe the hypermethylation of CHH and CHG at late stages, probably because their analysis relied on less time points during senescence progression. It is likely that discrepancies between the WGBS analysis in our study and that of Yuan et al. (2020) are partially due to differences in growth conditions, physiological states, bioinformatic analyses and sample sizes. We used three biological replicates comprising pooled samples from ∼ 20 plants in total, followed by a procedure based on a Bayesian hierarchical model utilized by the DSS package from bioconductor for DML and DMR scoring, whereas Yuan et al. (2020) used two biological replicates and Fischer’s exact test for DMR discovery, without explicit counting of DMLs. Furthermore, their general analysis compared methylation levels at 100-kb windows, whereas our analysis was based on individual cytosines. Finally, our experiment proceeded beyond the 50% yellowing of the leaves defined as their last time point and at that later stage (SM), we observed the highest degrees of CHH hypermethylation. Still, the proposed concerted methylation loss particularly in senescence-associated genes or their promoters, as well as the proposed delay of senescence by hypermethylation (Yuan et al., 2020) are not supported by our experimental setup.

Despite individual methylated cytosines may have profound effects on transcription factor binding to methylated DNA (O’Malley et al., 2016), only accumulations of differentially methylated cytosines in genomic regions are generally considered genetically relevant (Quadrana and Colot, 2016). Biochemical binding assays to methylated WRKY target sites strongly suggested that WRKYs ability to bind their target sites is strongly prohibited by methylation, but it is important to note that we used only fully methylated targets in our binding assays (Fig. 2G, Suppl. Fig. S3). While methylation within the chromosomal DNA of a cell is either present or not, gradually smaller methylation changes in a tissue suggest that target binding is affected in less cells, with limited effect on average gene expression within a tissue. Strong effects of methylation on target binding are supported by independent experiments (O’Malley et al., 2016).

To find genes potentially regulated via cytosine methylation, we searched for DMRs located closely to their transcription start sites (Fig. 5). Cytosine methylation around the TSS and in promoter regions have long been known to be involved in the regulation of gene expression (Finnegan et al., 1996; Jeddeloh et al., 1998; Wang et al., 2004). Interestingly, we have found a hypomethylated DMR lying in close proximity to the TSS of the demethylase gene *ROS1* (Fig. 6). Lei et al., (2015) have described a 39-bp DNA methylation monitoring system (MEMS) found between the TSS of *ROS1* and an adjacent helitron transposon element. Low methylation levels of this region are associated with the reduced expression of *ROS1*, whereas high methylation levels correlate with a higher expression (Lei et al., 2015; Fig. 6). The decline of *ROS1* expression with the progression of senescence suggests reduced demethylation activity and strengthens the idea that the loss of methylation during senescence is attributable to inhibited maintenance, rather than to active demethylation (Fig. 1, 2).

The cytosine methylation changes during progressing leaf senescence were not preferentially associated with any physiological gene class, e.g. with nutrient status, defence- or senescence-associated genes. For the key regulatory gene *WRKY53*, no changes in its cytosine methylation have been observed, despite its upregulation during early senescence (Zentgraf et al., 2010). Because high-resolution transcriptomics datasets of progressing senescence exist (e.g. Breeze et al., 2011; Yuan et al., 2020) we did not obtain another transcriptome dataset, but concentrated just on a few genes. We confirmed a CHH DMR in the 2000bp region adjacent to the *PHYC* gene (Fig. 6, Yuan et al., 2020) and a differentially methylated A-box in its promoter. Changes in *PHYC* expression were correlated with this DMR, but the temporal inspection of other senescence-associated genes such as those identified in Yuan et al., (2020) failed to support the simple conclusion that DMRs and senescence-associated gene expression were causally or temporally linked (Fig. 6, Yuan et al., 2020). Many other recent studies also have failed to identify a general, simple direct relationship between methylation changes in tissues and gene expression changes (Quadrana and Colot, 2016; Zhang et al., 2018). Other epigenetic factors, such as the link with chromatin (de-)condensation or histone marks, are probably involved.

Aberrant methylation in transcription factor binding sites may have caused disturbances in signal transduction networks during senescence, e.g. of the WRKY53, WRKY18 and WRKY25 interaction network (Zentgraf and Doll, 2019). WRKY transcription factors preferentially bind the *cis*-element *TTGACT/C*, named a W-box, that contains cytosines in CHH or, in some cases, CHG contexts. Full methylation of target sites impairs WRKY binding *in vitro* (Fig. 2; Suppl. Fig. S3). By an independent and different approach, O’Malley et al., (2016) have demonstrated that DNA binding of more than 75% of transcription factors in *Arabidopsis* is sensitive to methylation, including WRKY18 and WRKY25. Methylation in W-boxes (in the CHH context) changed between stages, this happened most strongly in TEs, followed by ORFs. W-boxes in regions adjacent to the TSS of ORFs were least methylated (Fig. 7). We speculate that cytosine methylation in genomic regions other than promoters serves to redirect transcription factors from unwanted non-functional sites. Therefore, the disruption of methylation maintenance in CHH and CHG contexts in the *ddc* mutant might indirectly affect the concentration of free unbound WRKY transcription factors and ultimately affect complex signalling networks. A link to aberrant WRKY binding in the *ros1* mutant is also supported by the finding that ROS1 erases methylation in certain promoters with WRKY binding sites (Halter et al., 2021). A-, C- and G-boxes also have the lowest methylation when in close proximity to transcription start sites, imposing limited access of methyltransferases to these regions, resulting in lower methylation levels. Any *Arabidopsis* gene has been estimated to be subjected to an average of 25-75 transcription factor binding events and most of these are expected to have little effect on gene expression (Jones and Vandepoele, 2020; Ferrier et al., 2011). As approximately 5% of genes in *A. thaliana* are methylated in promoter regions, we therefore expect for these a complex cumulative relationship of individual DMLs and DMRs with gene expression.

We present a schematic model summarizing the observed links of cytosine methylation with senescence progression and generative growth (Fig. 8). The quicker senescence progression and nitrogen remobilization in the *ddc* and *ros1* mutants is related to altered C:N ratios and may be secondary. The majority of cytosine methylation remains remarkably stable, although overall cytosine methylation transiently decreased during the initial phase (Fig. 4). The majority of hypermethylated loci at the ultimate was in CHH contexts, pointing to the involvement of functional RdDM and potentially of the DRM1 and DRM2 methyltransferases. Initial hypomethylation has been observed predominantly in CG contexts, indicating the inhibition of the maintenance of methylation primarily at earlier stages. The regulation of *ROS1* via differential methylation at its own promoter region early after flower induction (Fig. 6) indicates a possible downregulation of active demethylation, which is missing in *ros1* mutants. Despite that the DNA methylation pattern changes with senescence progression in the leaf we consider any direct effect on expression of key senescence-associated genes unlikely.

**Figure 8.**
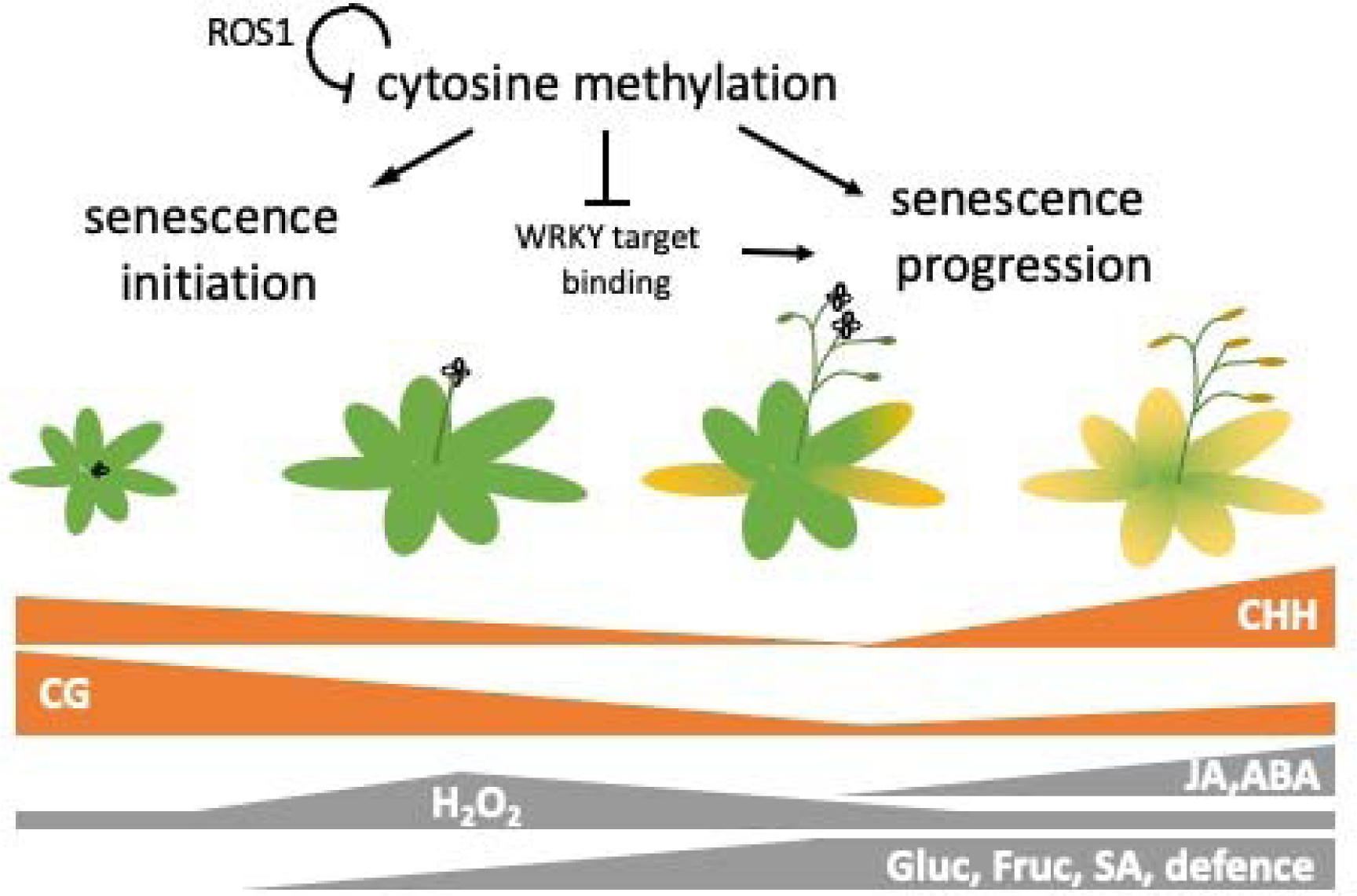
Relationship between flowering, senescence and methylation. Changes in physiological indicators and cytosine methylation during the senescence process at the four time points, namely Bolting, Flowering, Seed development and Seed maturation, are given. CHH, CG: context specific methylation; Gluc: glucose; Fruc: fructose; H_2_O_2_: hydrogen peroxide; ABA: abscisic acid, SA: salicylic acid; JA: jasmonic acid.

## Conclusions

Our results indicate that moderate methylation loss and the establishment of *de novo* methylation are associated with execution of the leaf senescence program and nitrogen remobilization, which is altered in methylation mutants.

## Abbreviations

CN ratio: carbon:nitrogen ratio
C: cytosine
G: guanine
H: any base of A,T,C
TSS: transcriptional start site
ORF: open reading frame
TF: transcription factor
ROS: reactive oxygen species
JA: jasmonic acid
SA: salicylic acid
ABA: abscisic acid
RdDM: RNA-directed
DNA: methylation
LD: long day
*ROS1*: repressor of silencing1
WGBS: whole genome bisulfite sequencing
ACA: automated colorimetric assay
DML: differentially methylated loci
DMRs: differentially methylated region
Blt: bolting
Flwr: Flowering
SD: seed development
SM: seed maturation
PCA: principle component analyses
TE: transposable element
DAS: days after sowing.

## Materials and methods

### Comparative analyses of *ddc, ros1* and Col-0

The hypermethylated *ros1* and hypomethylated *ddc* mutant lines, used previously by Chen et al. (2018), were grown in hydroponic culture under long days (16/8h day/night) and compared with Col-0 wild type for visual symptoms of senescence (for details, see **Supplementary Material**). Experiments were conducted and repeated in two different laboratories (Hohenheim and Tübingen) with similar outcomes. Flowering time was scored as the percentage of plants that showed the visible transition of the shoot apical meristem to reproductive growth for each scoring date. Images of the plants were taken every two to three days and were evaluated visually for symptoms of senescence. First symptoms of senescence were scored as the date when visible yellowing appeared on the leaf rosette. Fifty percent senescence was scored as the date when approximately half of the leaf rosette showed visible discoloration. Statistical analysis was performed in SAS by *glimmix* and *lsmeans*. The three genotypes were then grown together for 50 days under short day conditions (8/16h day/night), followed by a transition to long days. For Col-0, samples were taken at 30 DAS (day 0 after light change), 44 DAS (day 14), 51 DAS (day 21), 60 DAS (day 30) and 90 DAS (day 60). The last samples were taken when senescence was complete for the whole plant. The above-ground biomass was separated into leaf, stem and seed material and weighed separately. Statistical analysis was performed with R, interactions were analysed using the functions *aov*, *lm*, *lsmeans* and *cld,* with *alpha* = 0.05 and *adjust* = ‘tukey’, and the *LSD.test* was performed following *aov*. The three genotypes were compared separately for rosette, stem and seed biomass.

### Leaf physiology and cytosine methylation

Col-0 plants were grown in soil under long day conditions. During vegetative growth, the leaves at positions 5 to 9 within the rosette were labelled from bottom to top for the following analyses: leaf 5 – chlorophyll extraction; leaf 6 – sugar content; leaf 7 – whole genome bisulfite sequencing (WGBS); leaf 8 – hormone analyses; leaf 9 – H_2_O_2_ content. Four developmental stages were chosen roughly corresponding to the following stages from the “Timetable of *Arabidopsis* Growth Stages” in https://www.arabidopsis.org/: 1 – S5 Inflorescence emergence (Bolting: Blt); 2 –S6 Flower production (Flowering: Flwr); 3 – S6.3 30% of flowers produced open (Seed development: SD); 4 – S8 Silique or fruit ripening (Seed maturation: SM) (Figure 2). Twenty plants were harvested at each developmental stage for further analyses.

### Leaf physiology

Automated colorimetric assay (ACA), chlorophyll extraction and hydrogen peroxide levels (H_2_O_2_) were estimated as described by Bresson et al. (2018). Principle component analysis (PCA) was performed with R by using pixel counts from the ACA analysis for green, green-yellow, yellow, brown and total leaf pixels and by using leaf fresh weights with the function *prcomp*. Linear regression analysis with k-mer validation for the correlation of ACA with chlorophyll concentrations was performed with the ‘*ceret*’ package of R. Data indexing was performed using the *createMultiFolds* function with *k =* 5 and *times =* 10. The method was adjusted via *trainControl,* with *method* = ‘*repeatedcv’, number =* 5, *repeat =* 10 and *index*. The model was trained using *train,* with *method = ‘lm’*. Sugar content was analysed from leaf number 6 in a Dionex/Thermo ICS 5000 system equipped with pulsed amperiometric detection. Hormone levels were quantified from leaf number 8 by using GCMS (Shimadzu TQ8040) in the splitless MRM Mode.

### Cytosine methylation analyses

DNA extracted from leaf number 7 (triplicate DNA samples per timepoint, pooled from ∼20 leaves) was sequenced by Novogene Co. Ltd. on the Illumina 150 PE platform with a coverage of 3G (22x). The methylome raw dataset is available upon publication in the Gene Expression Omnibus (GEO) database under the accession number GSE176468. The quality of the raw reads was assessed using FastQC Version 0.11.9. Trimmomatic 0.39 was employed to trim and filter the raw reads. Sequence alignment was performed using Bismark 0.22.3 at default settings (Krueger and Andrews, 2011). The reference genome was TAIR10 assembly. Following analysis of the coverage distribution, cytosines with a coverage of <3 or >40 were removed from further analyses using R. Differential methylation was statistically analysed using the DSS package from bioconductor (Park and Wu, 2016). PCA and cluster analyses were performed on R by using the weighted means for methylation of 50 bp bins of the entire genome and with all single cytosines recognized as DMLs in any of the possible pairwise comparisons. Differential methylated loci (DML, at least 25% change in methylation) were called using the DSS package with the *callDML* function at a significance level of 0.05. Differential methylated regions were called using the *callDMR* function with *p.threshold* = 0.05.

The locations of transposon elements (TEs) and gene open reading frames (ORFs) were downloaded from the TAIR10 assembly and customized into the *bed* file format by using python. The region lying 2500bp upstream of the ATG start codon of every ORF was also recorded in the *bed* format and analysed as the 2500bp promoter. The *bedtools* functions *intersect* and *closest* were used to find the locations of all DML and DMRs with respect to the above-mentioned genomic elements. The locations of all W-boxes (*TTGACT/C*) were searched in the TAIR10 assembly with the regular expression function of python, namely *re.finditer*. Bootstrap analysis was performed in R to estimate the statistical significance of cytosine methylation in transcription factor binding sites compared with their genomic context.

### Gene expression analysis by qRT-PCR

RNA was extracted from the ground leaf powder left over after the WGBS experiment (leaf 7) by using Analytikjena innuPREP Plant RNA Kit with Guanidinium Chloride. The *AtACTIN2* gene was used as a reference gene for normalization. Six other genes were chosen for analysis: *AtROS1* (*AT2G36490*)*, AtPHYC* (*AT5G35840*), HISTONE CHAPERONE HIRA-like protein (*AT3G44530*), Peptidase M20/M25/M40 family protein (*AT1G44820*), *SPERMIDINE SYNTHASE 3* (*AT5G53120*) and 50S ribosomal-like protein (*AT1G66890*). The primers used are listed in Supplementary Table S1.

### Protein expression and DPI-ELISA with WRKY transcription factors

WRKY18, WRKY25 and WRKY53 were expressed in *E. coli* with an N-terminal fused 6xHis-tag, as described by Doll et al., (2020). DPI-ELISA was performed following the instructions from Brand et al., (2010). A 29-bp-long biotinylated artificial sequence containing 3 W-boxes was used for the assay. Detailed description of the methods used in this study are given in **Supplementary Materials**.

## Authors contribution

EV, UL and UZ designed the experiments, EV carried out the experiments and the computational analyses and prepared all the figures. EV wrote the first manuscript draft with the help of UL. UL and UZ revised the manuscript and contributed to the finalization of the manuscript.

## Acknowledgements

We thank Dr Ute Bertsche of the Core facility of the University of Hohenheim for cytokine analysis, Helene Ochott, Elke Dachtler, Charlotte Haake and Deborah Schnell for technical support and nutrient analyses, Dr. Jasmin Doll, ZMBP, University of Tübingen for her help with the DPI-ELISA and the cytosine methylation experiments, Dr. Stephan Bieker, ZMBP, University of Tübingen for his help with R, Dr. Mark Stahl, Bettina Stadelhofer and Dr. Joachim Kilian, of the central facilities at the ZMBP for help with sugar and hormone analytics and R. Theresa Jones for editorial comments on the manuscript.

## Legends of supplemental figures and tables

**Suppl. Figure S1.** Visual appearance of the leaf rosette at transition to flowering. From left to right: Col-0, *ddc* and *ros1.* Note that *ros1,* but not the other two genotypes, already has one senescent leaf.

**Suppl. Figure S2.** Analyses of cytosine methylation data of *ddc* and Col-0 based on methylome data from Stroud et al., (2013). (A) Cytosine methylation levels in CG, CHG and CHH contexts. (B) Number of DMRs (out of a total of 228) intersecting with ORFs, 2500bp promoter regions and TEs.

**Suppl. Figure S3.** Influence of cytosine methylation on the binding of WRKY25 and WRKY18 via DPI-ELISA. (A) Affinity of WRKY18 to an artificial promoter sequence with or without cytosine methylation. 0.2 and 0.02 Art indicate 20 pmol and 2 pmol of double-stranded artificial promoter DNA fragments per 60 µl, respectively. 5mC indicates methylated DNA. (B) Affinity of WRKY25 to an artificial promoter sequence with (5mC) or without cytosine methylation. Colour codes indicate the concentrations of protein (conc) used in µg per 60 µl reaction. (C) Double-stranded sequence of the artificial (Art) methylated promoter (5mC). n = 2; std. error; Each experiment was carried out with 2 technical replicates and was repeated at least twice.

**Suppl. Figure S4.** Four developmental stages were analysed during the experiment: (A) S 5 Bolting, (B) S 6 Flowering, (C) S 6.3 Seed development and (D) S 8 Seed maturation.

**Suppl. Figure S5.** Visual quantification of senescence in leaves numbered 5 to 9 from Col-0 plants. Automated colorimetric assay (ACA) was performed on pixel counts for green, green-yellow, yellow and brown coloration, total pixel count and leaf fresh weight (n = 20 ± SEM). (A) Leaf 5. (B) Leaf 6. (C) Leaf 7. (D) Leaf 8. (E) Leaf 9. (F) Chlorophyll in leaf 5 (n = 20 ± SD). (G) PCA for distinguishing the colours of the various leaves. (H) Glucose and fructose per predicted chlorophyll concentrations. (I) Hydrogen peroxide vs. chlorophyll concentrations plotted against normalized leaf weight. w.norm = normalized leaf weight on a scale of -1 to 1, where -1 to 0 indicates leaf growth and 0 to 1 indicates leaf desiccation and loss of weight. Chl.DW = predicted chlorophyll concentration in leaf dry weight.

**Suppl. Figure S6.** Change of defence-related secondary metabolites in leaf number 7. Various anthocyanins, namely calabricoside A, cyanidin and kaempferol, at four time points measured in arbitrary units (A.U.; n = 3; means ± SD).

**Suppl. Figure S7.** Distribution of differentially methylated cytosines in CG, CHG and CHH contexts along chromosome 1 in three pairwise comparisons.

**Suppl. Table S1.** Primers used.

**Suppl. Table S2.** Statistics of methylome sequencing.

**Suppl. Table S3.** Gene list of ORFs intersecting with DMRs.

**Suppl. Table S4.** Gene list of 2500 promoters intersecting with DMRs.

**Suppl. Table S5.** Genes/promoters with differentially methylated A-, C- or G-boxes.

**Suppl. Table S6.** Genes/promoters with differentially methylated W-boxes.

## References

Agüera E, De la Haba P (2018) Leaf senescence in response to elevated atmospheric CO 2 concentration and low nitrogen supply. Biol Plant 62: 401–408

Ay N, Irmler K, Fischer A, Uhlemann R, Reuter G, Humbeck K (2009) Epigenetic programming via histone methylation at WRKY53 controls leaf senescence in *Arabidopsis thaliana*. Plant J 5: 333–346

Beemster GT, De Veylder L, Vercruysse S, West G, Rombaut D, Van Hummelen P, Galichet A, Gruissem W, Inze D, Vuylsteke M (2005) Genome-wide analysis of gene expression profiles associated with cell cycle transitions in growing organs of *Arabidopsis*. Plant Physiol 138: 734–743

Brand LH, Kirchler T, Hummel S, Chaban C, Wanke D (2010) DPI-ELISA: a fast and versatile method to specify the binding of plant transcription factors to DNA in vitro. Plant Methods 6: 25

Breeze E, Harrison E, McHattie S, Hughes L, Hickman R, Hill C, Kiddle S, Kim Y, Penfold CA, Jenkins D, Zhang C, Morris K, Jenner C, Jackson S, Thomas B, Tabrett A, Legaie R, Moore JD, Wild DL, Ott S, Rand D, Beynon J, Denby K, Mead A, Buchanan-Wollaston V (2011) High-resolution temporal profiling of transcripts during *Arabidopsis* leaf senescence reveals a distinct chronology of processes and regulation. Plant Cell 23: 873–894

Bresson J, Bieker S, Riester L, Doll J, Zentgraf U (2018) A guideline for leaf senescence analyses: from quantification to physiological and molecular investigations. J Exp Bot 69: 769–786

Brusslan JA, Alvarez-Canterbury AMR, Nair NU, Rice JC, Hitchler MJ, Pellegrini M (2012) Genome-wide evaluation of histone methylation changes associated with leaf senescence in *Arabidopsis*. PLoS One 7: e33151

Buchanan-Wollaston V, Page T, Harrison E, Breeze E, Lim PO, Nam HG, Lin JF, Wu SH, Swidzinski J, Ishizaki K, Leaver CJ (2005) Comparative transcriptome analysis reveals significant differences in gene expression and signalling pathways between developmental and dark/starvation-induced senescence in *Arabidopsis*. Plant J 42: 567–585

Cao X, Jacobsen SE (2002) Locus-specific control of asymmetric and CpNpG methylation by th*DRM* and *CMT3* methyltransferase genes. Proc Nat Acad Sci USA 99: 16491–16498

Chan SWL, Henderson IR, Zhang X, Shah G, Chien JSC, Jacobsen SE (2006). RNAi, DRD1, and histone methylation actively target developmentally important non-CG DNA methylation in *Arabidopsis*. PLoS genetics 2: e83

Chen Q, Xu X, Xu D, Zhang H, Zhang C, Li G (2019) WRKY18 and WRKY53 coordinate with HISTONE ACETYLTRANSFERASE1 to regulate rapid responses to sugar. Plant Physiol 180: 2212–2226

Chen X, Lu L, Mayer KS, Scalf M, Qian S, Lomax A, Smith LM, Zhong X (2016) POWERDRESS interacts with HISTONE DEACETYLASE 9 to promote ageing in *Arabidopsis*. Elife 5: e17214

Chen X, Schönberger B, Menz J, Ludewig U (2018) Plasticity of DNA methylation and gene expression under zinc deficiency in *Arabidopsis* roots. Plant Cell Physiol 59: 1790–1802

Diaz C, Lemaître T, Christ A, Azzopardi M, Kato Y, Sato F, Morot-Gaudry JF, Le Dily F, Masclaux-Daubresse C (2008) Nitrogen recycling and remobilization are differentially controlled by leaf senescence and development stage in *Arabidopsis* under low nitrogen nutrition. Plant Physiol 147: 1437–1449

Doll J, Muth M, Riester L, Nebel S, Bresson J, Lee HC, Zentgraf U (2020) *Arabidopsis thaliana* WRKY25 transcription factor mediates oxidative stress tolerance and regulates senescence in a redox-dependent manner. Front Plant Sci 10: 1734

Dong J, Chen C, Chen Z (2003) Expression profiles of the *Arabidopsis* WRKY gene superfamily during plant defense response. Plant Mol Biol 51: 21–37

Dowen RH, Pelizzola M, Schmitz RJ, Lister R, Dowen JM, Nery JR, Dixon JE, Ecker, JR (2012). Widespread dynamic DNA methylation in response to biotic stress. Proc Nat Acad Sci USA 109: E2183–E2191

Dubrovina AS, Kiselev KV (2016) Age□associated alterations in the somatic mutation and DNA methylation levels in plants. Plant Biol 18:185–196

Ferrier T, Matus JT, Jin J, Riechmann JL (2011) *Arabidopsis* paves the way: genomic and network analyses in crops. Curr Opin Biotech 22: 260–270

Finnegan EJ, Peacock WJ, Dennis ES (1996) Reduced DNA methylation in *Arabidopsis thaliana* results in abnormal plant development. Proc Nat Acad Sci USA 93: 8449-8454

Finnegan EJ, Kovac KA, Jaligot E, Sheldon CC, Peacock JW, Dennis ES (2005) The downregulation of FLOWERING LOCUS C (FLC) expression in plants with low levels of DNA methylation and by vernalization occurs by distinct mechanisms. Plant J 44: 420–432

Forgione I, Wołoszy ska M, Pacenza M, Chiappetta A, Greco M, Araniti F, Abenavoli MR, Van Lijsebettens M, Bitońnti MB, Bruno L (2019) Hypomethylated drm1 drm2 cmt3 mutant phenotype of *Arabidopsis thaliana* is related to auxin pathway impairment. Plant Sci 280: 383–396

Gong Z, Morales-Ruiz T, Ariza RR, Roldan-Arjona T, David L, Zhu JK (2002) ROS1, a repressor of transcriptional gene silencing in *Arabidopsis*, encodes a DNA glycosylase/lyase. Cell 111: 803–814

Halter T, Wang J, Amesefe D, Lastrucci E, Charvin M, Rastogi MS, Navarro L (2021) The *Arabidopsis* active demethylase ROS1 cis-regulates defence genes by erasing DNA methylation at promoter-regulatory regions. Elife 10: e62994

Havé M, Marmagne A, Chardon F, Masclaux-Daubresse C (2017) Nitrogen remobilization during leaf senescence: lessons from *Arabidopsis* to crops. J Exp Bot 68: 2513–2529

He L, Wu W, Zinta G, Yang L, Wang D, Liu R, Zhang H, Zheng Z, Huang H, Zhang Q, Zhu JK (2018) A naturally occurring epiallele associates with leaf senescence and local climate adaptation in *Arabidopsis* accessions. Nat Commun 9: 1–11

Hickman R, Hill C, Penfold CA, Breeze E, Bowden L, Moore JD, … Buchanan-Wollaston V (2013) A local regulatory network around three NAC transcription factors in stress responses and senescence in *Arabidopsis* leaves. Plant J 75: 26–39

Ikeda Y, Kinoshita T (2009) DNA demethylation: a lesson from the garden. Chromosoma 118: 37–41

Jeddeloh JA, Bender J, Richards EJ (1998) The DNA methylation locus DDM1 is required for maintenance of gene silencing in *Arabidopsis*. Genes Dev 12: 1714–1725

Jibran R, Hunter DA, Dijkwel PP (2013) Hormonal regulation of leaf senescence through integration of developmental and stress signals. Plant Mol Biol 82: 547–561

Jones DM, Vandepoele K (2020) Identification and evolution of gene regulatory networks: Insights from comparative studies in plants. Curr Opin Plant Biol 54: 42–48

Jones MJ, Goodman SJ, Kobor MS (2015) DNA methylation and healthy human aging. Aging Cell 1: 924–932

Kim JI, Murphy AS, Baek D, Lee SW, Yun DJ, Bressan RA, Narasimhan ML (2011) YUCCA6 over-expression demonstrates auxin function in delaying leaf senescence in *Arabidopsis thaliana*. J Exp Bot 62: 3981–3992

Kinoshita Y, Saze H, Kinoshita T, Miura A, Soppe WJ, Koornneef M, Kakutani T (2007) Control of FWA gene silencing in *Arabidopsis thaliana* by SINE related direct repeats. Plant J 49: 38–45

Kou HP, Li Y, Song XX, Ou XF, Xing SC, Ma J, … Liu B (2011) Heritable alteration in DNA methylation induced by nitrogen-deficiency stress accompanies enhanced tolerance by progenies to the stress in rice (*Oryza sativa* L.). J Plant Physiol 168: 1685–1693

Krueger F, Andrews SR (2011) Bismark: a flexible aligner and methylation caller for Bisulfite-Seq applications. Bioinformatics 27: 1571–1572

Kuhlmann M, Meyer RC, Jia Z, Klose D, Krieg LM, von Wirén N, Altmann T (2020) Epigenetic variation at a genomic locus affecting biomass accumulation under low nitrogen in *Arabidopsis thaliana*. Agronomy 10: 636

Lang Z, Wang Y, Tang K, Tang D, Datsenka T, Cheng J, Zhang Y, Handa AK, Zhu JK (2017) Critical roles of DNA demethylation in the activation of ripening-induced genes and inhibition of ripening-repressed genes in tomato fruit. Proc Nat Acad Sci USA 114: E4511–E4519

Lei M, Zhang H, Julian R, Tang K, Xie S, Zhu JK (2015) Regulatory link between DNA methylation and active demethylation in *Arabidopsis*. Proc Nat Acad Sci USA 112: 3553–3557

Liebsch D, Keech O (2016) Dark induced leaf senescence: new insights into a complex light dependent regulatory pathway. Ne^LJ^w Phytol 212: 563–570

Lin YL, Tsay YF (2017) Influence of differing nitrate and nitrogen availability on flowering control in *Arabidopsis*. J Exp Bot 68: 2603–2609

Mager S, Ludewig U (2018) Massive loss of DNA methylation in nitrogen-, but not in phosphorus-deficient *Zea mays* roots is poorly correlated with gene expression differences. Front Plant Sci 9: 497

Masclaux-Daubresse C, Daniel-Vedele F, Dechorgnat J, Chardon F, Gaufichon L, Suzuki A (2010) Nitrogen uptake, assimilation and remobilization in plants: challenges for sustainable and productive agriculture. Ann Bot 105: 1141–1157

Matzke MA, Kanno T, Matzke AJ (2015) RNA-directed DNA methylation: the evolution of a complex epigenetic pathway in flowering plants. Ann Rev Plant Biol 66: 243–267

Ogneva ZV, Dubrovina AS, Kiselev KV (2016) Age-associated alterations in DNA methylation and expression of methyltransferase and demethylase genes in *Arabidopsis thaliana*. Biol Plant 60: 628–634

O’Malley RC, Huang SSC, Song L, Lewsey MG, Bartlett A, Nery JR, Galli M, Gallavotti A, Ecker JR (2016) Cistrome and epicistrome features shape the regulatory DNA landscape. Cell 165: 1280–1292

Park Y, Wu H (2016) Differential methylation analysis for BS-seq data under general experimental design. Bioinformatics 32: 1446–1453

Quadrana L, Colot V (2016) Plant transgenerational epigenetics. Ann Rev Genet 50: 467–491

Schippers, JH, Jing, HC, Hille, J, Dijkwel, PP (2007) Developmental and hormonal control of leaf senescence. Senescence processes in plants, 145–170. Oxford Blackwell Publishing

Sequeira-Mendes J, Aragüez I, Peiró R, Mendez-Giraldez R, Zhang X, Jacobsen SE, Bastolla U, Gutierrez C (2014) The functional topography of the *Arabidopsis* genome is organized in a reduced number of linear motifs of chromatin states. Plant Cell 26: 2351–2366

Stroud H, Greenberg MV, Feng S, Bernatavichute YV, Jacobsen SE (2013) Comprehensive analysis of silencing mutants reveals complex regulation of the *Arabidopsis* methylome. Cell 152: 352–364

Takasaki H, Maruyama K, Takahashi F, Fujita M, Yoshida T, Nakashima K, … Shinozaki K (2015) SNAC-As, stress-responsive NAC transcription factors, mediate ABA-inducible leaf senescence. Plant J 84: 1114–1123

Trejo-Arellano MS, Mehdi S, de Jonge J, Tomastíková ED, Köhler C, Hennig L (2020) Dark-induced senescence causes localized changes in DNA methylation. Plant Physiol 182: 949–961

Wang J, Tian L, Madlung A, Lee HS, Chen M, Lee JJ, Watson B, Kagochi T, Comai L, Chen ZJ (2004) Stochastic and epigenetic changes of gene expression in *Arabidopsis* polyploids. Genetics 167: 1961–1973

Wingler A, Purdy S, MacLean JA, Pourtau N (2006) The role of sugars in integrating environmental signals during the regulation of leaf senescence. J Exp Bot 57: 391–399

Yong-Villalobos L, González-Morales SI, Wrobel K, Gutiérrez-Alanis D, Cervantes-Peréz SA, Hayano-Kanashiro C, … Herrera-Estrella L (2015) Methylome analysis reveals an important role for epigenetic changes in the regulation of the *Arabidopsis* response to phosphate starvation. Proc Nat Acad Sci USA 112: E7293–E7302

Yuan L, Wang, D, Cao L, Yu N, Liu K, Guo Y, Gan S, Chen L (2020) Regulation of leaf longevity by DML3-mediated DNA demethylation. Mol Plant 13: 1149–1161

Zentgraf U, Doll J (2019) *Arabidopsis* WRKY53, a node of multi-layer regulation in the network of senescence. Plants 8: 578

Zentgraf U, Laun T, Miao Y (2010) The complex regulation of WRKY53 during leaf senescence of *Arabidopsis thaliana*. Eur J Cell Biol 89: 133–137

Zhang H, Lang Z, Zhu JK (2018) Dynamics and function of DNA methylation in plants. Nat Rev Mol Cell Biol 19: 489–506

Zicola J, Liu L, Tänzler P, Turck F (2019) Targeted DNA methylation represses two enhancers of FLOWERING LOCUS T in *Arabidopsis thaliana*. Nat Plant 5: 300–307

