## Supplementary material for "Moderate DNA methylation changes associated with nitrogen remobilization and leaf senescence in *Arabidopsis*": Suppl Figs and Methods

**Suppl. dataset 1 (Suppl. figures with legends and suppl. Methods)**

For: Control of nitrogen remobilization and senescence in *Arabidopsis thaliana* by DNA methylation dynamics

By Emil Vatov et al.

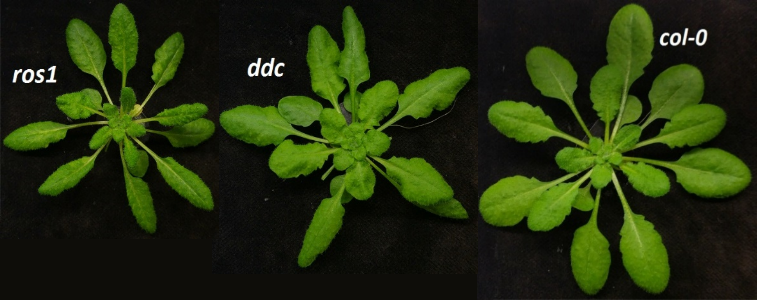

**Suppl. Figure S1.** Visual appearance of the leaf rosette at transition to flowering. From left to right: Col-0, *ddc* and *ros1.* Note that *ros1,* but not the other two genotypes, already has one senescent leaf.

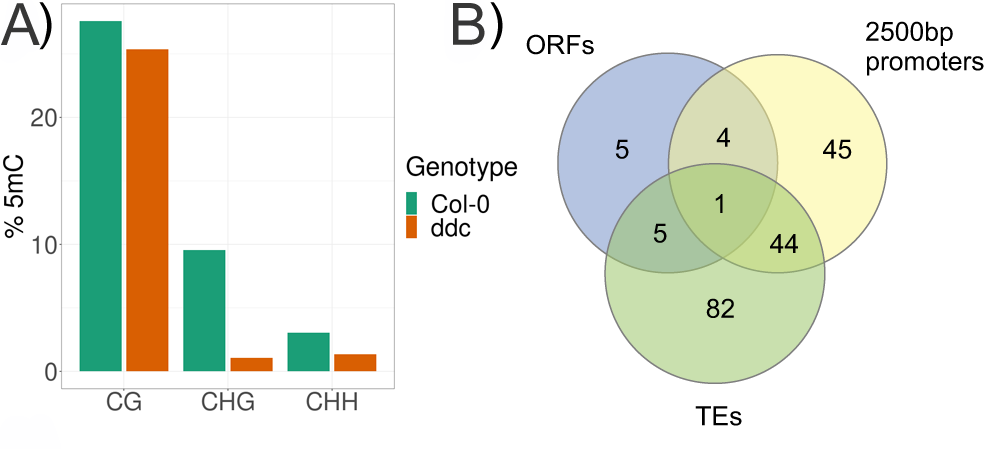

**Suppl. Figure S2.** Analyses of cytosine methylation data of *ddc* and Col-0 based on methylome data from Stroud et al., (2013). (A) Cytosine methylation levels in CG, CHG and CHH contexts. (B) Number of DMRs (out of a total of 228) intersecting with ORFs, 2500bp promoter regions and TEs.

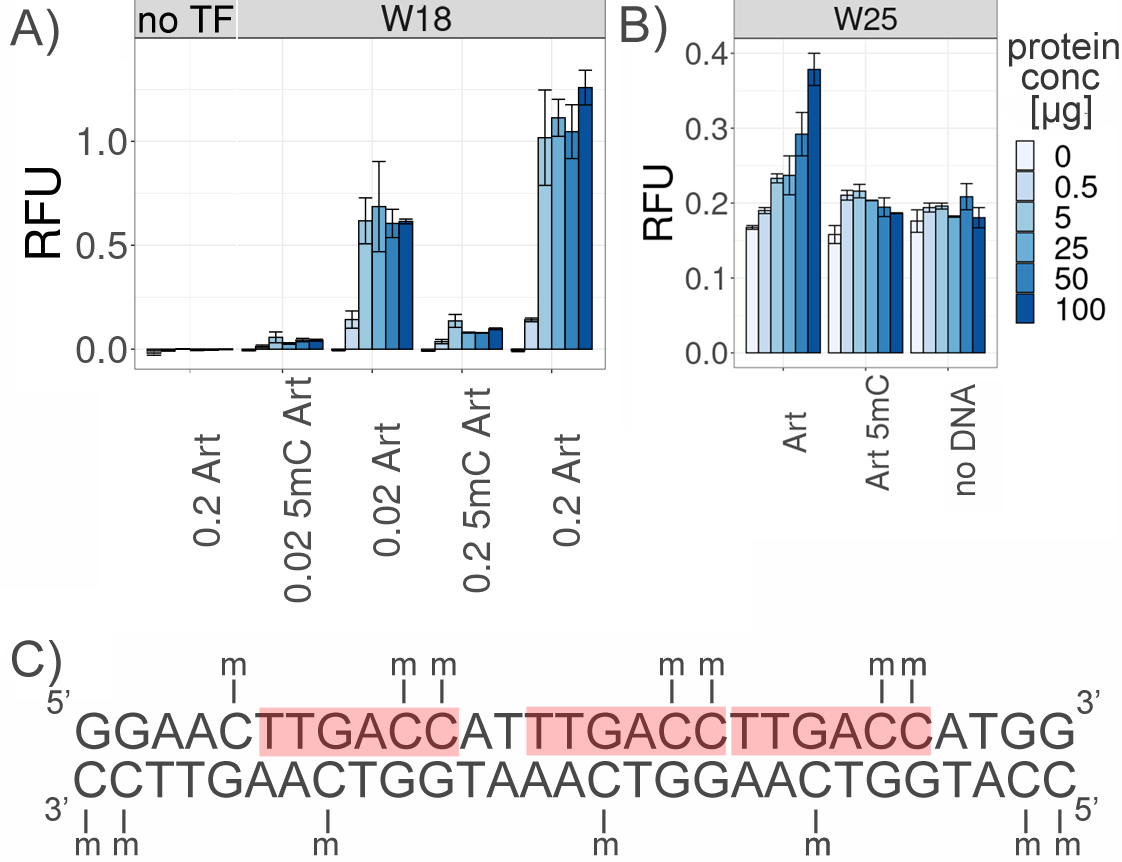

**Suppl. Figure S3.** Influence of cytosine methylation on the binding of WRKY25 and WRKY18 via DPI-ELISA. (A) Affinity of WRKY18 to an artificial promoter sequence with or without cytosine methylation. 0.2 and 0.02 Art indicate 20 pmol and 2 pmol of double-stranded artificial promoter DNA fragments per 60 µl, respectively. 5mC indicates methylated DNA. (B) Affinity of WRKY25 to an artificial promoter sequence with (5mC) or without cytosine methylation. Colour codes indicate the concentrations of protein (conc) used in µg per 60 µl reaction. (C) Double-stranded sequence of the artificial (Art) methylated promoter (5mC). W-boxes are highlighted in red colour. Each experiment was carried out with 2 technical replicates and was repeated at least twice.

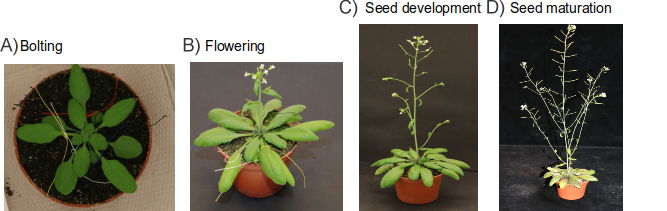

**Suppl. Figure S4.** Four developmental stages were analysed during the experiment: (A) Bolting, (B) Flowering, (C) Seed development and (D) Seed maturation.

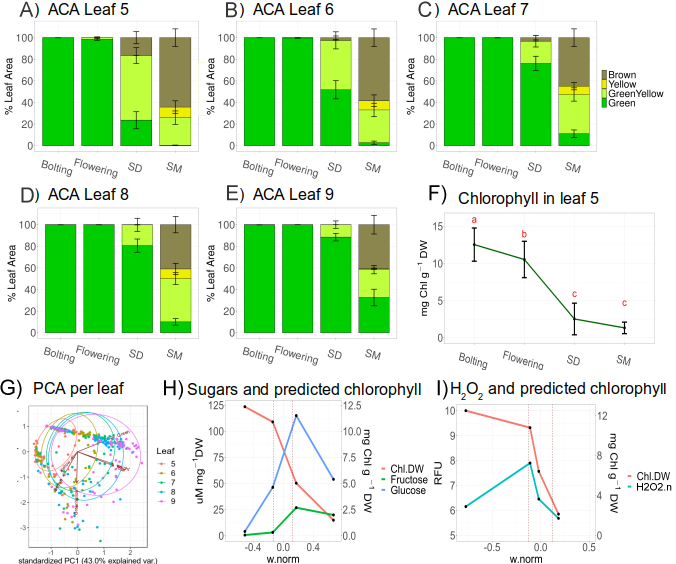

**Suppl. Figure S5**. Visual quantification of senescence in leaves numbered 5 to 9 of Col-0 plants. Automated colorimetric assay (ACA) was performed on pixel counts for green, green-yellow, yellow and brown coloration, total pixel count and leaf fresh weight (n = 20 ± SEM). (A) Leaf 5. (B) Leaf 6. (C) Leaf 7. (D) Leaf 8. (E) Leaf 9. (F) Chlorophyll in leaf 5 (n = 20 ± SD). (G) PCA for distinguishing the colours of the various leaves. (H) Glucose and fructose per predicted chlorophyll concentrations. (I) Hydrogen peroxide vs. chlorophyll concentrations plotted against normalized leaf weight. w.norm = normalized leaf weight on a scale of -1 to 1, where -1 to 0 indicates leaf growth and 0 to 1 indicates leaf desiccation and loss of weight. Chl.DW = predicted chlorophyll concentration in leaf dry weight.

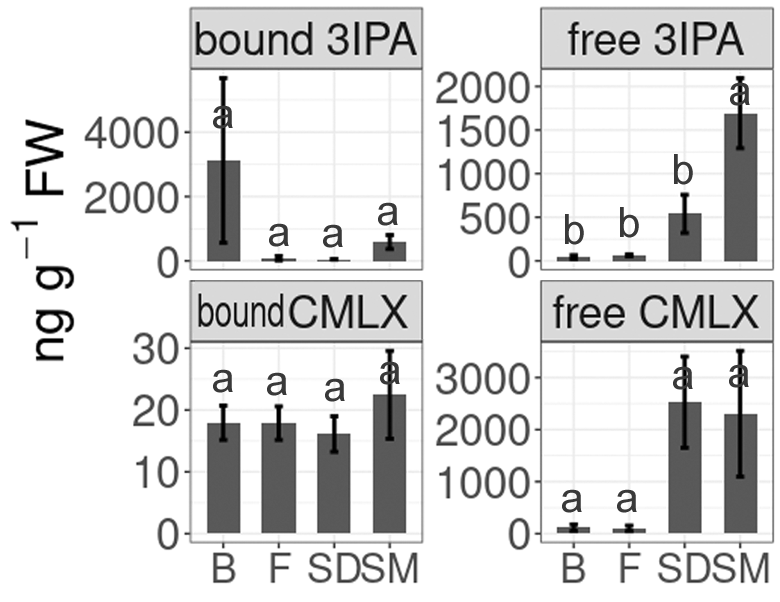

**Suppl. Figure S6.** Change of auxin precursor 3IPA and defence-related phytoalexins in bound and free form in leaf number 7, at four time points measured in arbitrary units (A.U.; n = 3; means ± SD).

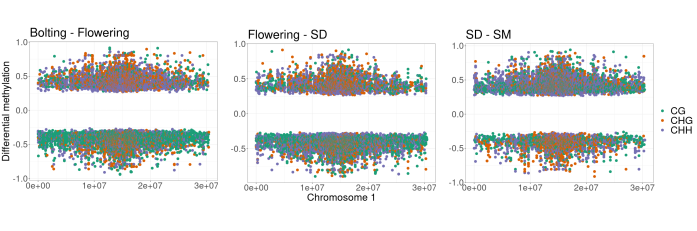

**Suppl. Figure S7.** Distribution of differentially methylated cytosines in CG, CHG and CHH contexts along chromosome 1 in three pairwise comparisons.

**Suppl. Materials and methods**

**Visual symptoms of senescence**

*ros1* hypermethylated and *ddc* hypomethylated mutant lines, earlier used by Chen et al., (2018), were grown in hydroponic culture under long days (16/8h day/night) and compared with Col-0 wild type for visual symptoms of senescence. After seeding, the seeds were stratified at 4^o^C for 5 days in the dark. ¼ Modified Hoagland was used, containing 1mM NH_4_NO_3_, 1mM CaCl_2_, 0.5mM MgSO_4_, 1mM KH_2_PO_4_, 1mM KNO_3_, 100µM NaFe(III)EDTA, 46µM H_3_BO_3_, 9µM MnSO_4_, 0.765µM ZnSO_4_, 0.32µM CuSO_4_, 0.016µM Na_2_MoO_3_. The air temperature was set at 20^o^C during the day and 16^o^C during the night. Photon intensity was 120-140 micro molar per square meter and humidity was set to 55% RH. 16 five-litre pots with 5 plants per pot were grown in a randomized complete block design. At least one plant per genotype was present in each pot. Flowering time was scored as the percentage of plants that showed visible transition of the shoot apical meristem to reproductive growth for each scoring date. Images of the plants were taken every two to three days and evaluated visually for symptoms of senescence. Statistical analysis was performed on SAS by using *glimmix* and *lsmeans*.

**Nitrogen remobilization during senescence**

The three genotypes were grown in 20 five-litre containers, as described for the previous experiment. The P concentration in the ¼ Modified Hoagland used previously was noticed to have an inhibiting effect on plant growth. Here, KH_2_PO_4_ was reduced to 0.2mM with the addition of 0.8mM KCl to compensate for K nutrition. Plants were grown for 50 days under short day illumination, followed by a transition to long day illumination for flower induction. Samples were taken starting at 30 DAS (day 0) and continuing at 44 DAS (day 14), 51 DAS (day 21), 60 DAS (day 30) and 90 DAS (day 60). The last samples were taken when senescence was complete for the whole plant. The above-ground biomass was separated into leaf, stem and seed and weighed separately. Statistical analysis was carried out with R, interactions were analysed using the functions *aov*, *lm*, *lsmeans* and *cld,* with *alpha* = 0.05 and *adjust* = ‘tukey’, and the *LSD.test* was performed following *aov*. The three genotypes for rosette, stem and seed biomass were compared separately.

**Leaf physiology and cytosine methylation during senescence**

120 Col-0 plants were grown in soil under long day conditions. During vegetative growth, the 5^th^ to 9^th^ leaves were labelled from bottom to top of the rosette for the following analyses: 5 – chlorophyll extraction; 6 – sugar content; 7 – whole genome bisulfite sequencing (WGBS); 8 – hormone analyses; 9 – H_2_O_2_ content. Four developmental stages were chosen relative to senescence, roughly corresponding to the following stages from the Timetable of Arabidopsis Growth Stages in <https://www.arabidopsis.org/>: 1 – S 5 Inflorescence emergence (Bolting); 2 –S 6 Flower production (Flowering); 3 – S 6.3 30% of flowers produced to be opened (Seed development: SD); 4 – S 8 Silique or fruit ripening (Seed maturation: SM) (Figure 1). 20 plants were harvested at each developmental stage for further analyses.

**Automated colorimetric assay (ACA) and chlorophyll extraction**

During harvest, leaves numbered 5-9 were separated and photographed for the automated colorimetric assay, as published earlier by Bresson et al., (2018). Each leaf was selected separately on ImageJ. To produce the ratios of green, yellow-green, yellow, brown and purple coloration for each leaf, pixel analysis of the images was performed in R. To assess the reliability of this assay, we extracted the chlorophyll of leaves numbered 5 and compared the results.

Chlorophyll was extracted as described by Bresson et al., (2018) with slight modifications, from 20 leaves per developmental stage. The fresh weight of each leaf was measured at harvest. The leaves were subsequently frozen in liquid nitrogen and then vacuum-dried. Once the dry weights had been measured, chlorophyll extraction in 80% acetone in phosphate buffer proceeded exactly as described in Bresson et al., (2018).

PCA analysis was performed in R by using the *prcomp* function with *center =* T and *scale =* T and by using pixel counts from the ACA analysis for green, green-yellow, yellow, brown and total leaf pixels and on leaf fresh weights.

Linear regression analysis with k-mer validation was performed using the ‘*ceret*’ package of R. Data was indexed using the *createMultiFolds* function with *k =* 5 and *times =* 10. The method was adjusted using *trainControl,* with *method* = ‘*repeatedcv’, number =* 5, *repeat =* 10 and *index*. The model was trained using *train,* with *method = ‘lm’*. The chlorophyll concentration in dry weight was predicted with the following formula:

*Chl.DW ~ percent.green + percent.green:g.norm + percent.green:percent.gy*

Chl.DW – mg chlorophyll per g dry weight

percent.green – Green pixel count / Total pixel count

percent.gy – GreenYellow pixel count / Total pixel count

g.norm – Normalization of green pixel count for each individual leaf towards the maximum green pixel count reached by each leaf position

(Maximum green pixel count – Leaf green pixel count)/Maximum green pixel count

**Sugars**

Sugar content was analysed in leaf number 6. Five leaves were pooled together at random. The sugar content was analysed in leaf number 6 by using a Dionex/Thermo ICS 5000 system equipped with pulsed amperiometric detection. Five leaves were pooled together at random to produce 4 replicates per developmental stage. Freeze-dried samples were homogenized with a ball mill and afterwards extracted with 500 µl of 80 % methanol, followed by a second extraction step with 500 µl of 20 % methanol (with 0.1 % formic acid). Both supernatants were combined and dried down in a vacuum concentrator. The samples were redissolved in 200 µl water and analysed on a CarboPac PA20 column from Thermo. Integration of the carbohydrate peaks and external calibration were used for quantification.

**Whole genome bisulfite sequencing**

DNA was extracted from leaf number 7 for whole genome bisulfite sequencing. The leaves were flash-frozen in liquid nitrogen and stored at -80^o^C. Six leaves were pooled together at random to produce 3 replicates per developmental stage. The pooled leaves were ground together in liquid nitrogen to give a fine powder. Approximately 100 mg plant material was added to 800 µl SLS Buffer + 40 µl Proteinase K (analytikjena Innu prep plant DNA kit) and incubated at 65^o^C for 1h. The samples were loaded on a prefilter (analytikjena Innu prep plant DNA kit) and centrifuged at 11,000x g for 1 min. An aliquot of 8 µl of 100 ng RNAseA/µl was added to each sample, which was then incubated at 37^o^C for 30 min. DNA was separated in 800 µl phenol:chloroform:isoamylalcohol (25:24:1) followed by centrifugation for 20 min at 4^o^C at maximum speed (14000 g). DNA was precipitated in 700 µl ice cold isopropanol with 70 µl 3M Na Acetate (pH 4.8) following incubation at -25^o^C for 40 mins. To wash the DNA pellet, 750 µl of 70% ethanol was added to the DNA and the mixture was incubated at room temperature for 20 min. The washing step with ethanol was repeated 3 times. The DNA pellet was dissolved in 50 µl of TE buffer, pH 8.

The DNA samples were sent to Novogene Co. Ltd , for WGBS on the Illumina, 150 PE platform with a coverage of 3G (22x). A total amount of 5.2 µg genomic DNA spiked with 26 ng lambda DNA was fragmented by sonication to lengths of 200-400 bp with Covaris S220, followed by end repair and adenylation. Cytosine-methylated barcodes were ligated to sonicated DNA as recommended by the manufacturer. The DNA fragments were then treated twice with bisulfite by using the EZ DNA Methylation-Gold^TM^ Kit (Zymo Research). The resulting single-stranded DNA fragments were PCR-amplified using KAPA HiFi HotStart Uracil + ReadyMix (2X). The library concentration was quantified by Qubit® 2.0 Flurometer (Life Technologies, CA, USA) and quantitative PCR, and the insert size was checked on the Agilent 2100 system. The clustering of the index-coded samples was performed on a cBot Cluster Generation System by using PE Cluster Kit cBot-HS (Illumina) according to the manufacturer’s instructions. Following cluster generation, the library preparations were sequenced on an Illumina platform and paired-end reads were generated. Image analysis and base calling were performed with the standard Illumina pipeline and, finally, paired-end reads were generated.

**qRT-PCR**

We tested the gene expression of six genes that were selected for further analysis. RNA was extracted from the ground leaf powder remaining after the WGBS experiment (leaf number 7) by using the Analytikjena innuPREP Plant RNA Kit with Guanidinium Chloride. RNA quantity and quality were tested via NanoDrop 2000c and gel electrophoresis. cDNA was synthesized from 1 µg of the total extracted RNA by using the Quantitec Revers transcription kit from Qiagen. qRT-PCR was performed with 15 ng cDNA per reaction in a Bio-rad CFX 384 with the GreenMasterMix from Genaxxon bioscience, according to the manufacturer’s protocol. PCR cycling parameters were set as follows: 95°C for 3 min, 45 cycles of 3 s at 95°C, 20 s at 60°C, and a final melting curve of 65 to 95°C with increments of 0.2°C.

The *AtACTIN2* gene was used as a reference gene for normalization. Six other genes were chosen for analysis: *AtROS1* (AT2G36490)*, AtPHYC* (AT5G35840), AT3G44530, AT1G44820, AT5G53120 and AT1G66890. The primers used are listed in Supplementary Table 1. Because of the limited amount of sample material, qRT-PCR was performed from two biological replicates with 3 technical repetitions. The experiment was repeated until consistent results were achieved. Insufficient RNA was obtained in the extract from the last stage of the experiment ‘SM’ (Seed Maturation) and, hence, the results presented here are from only one biological replicate. ΔΔCt values were calculated for each biological replicate. The timepoint ‘Flwr’ (Flowering) was taken as the reference; therefore, its mean value always equals 1 after normalization.

For each gene and technical replicate:

ΔCt = Ct_Blt_ – mean(Ct_Flwr_)

mean(Ct_Flwr_) is the average Ct from all technical replicates

For each gene, technical replicate and timepoint:

ΔΔCt = ΔCt_Gene_ - mean(ΔCt_actin_)

mean(ΔCt_actin_) is the average ΔCt from all technical replicates

ANOVA followed by TukeyHSD was performed in R to assess statistical significance as follows:

aov(2^-ΔΔCt^ ~ Bio + Timepoint)

This function performs an ANOVA for the effects of Bio (Biological replicate) and Timepoint on the normalized ΔΔCt value.

HSD.test(model,'Timepoint') provides the letter description following the TukeyHSD test for statistical significance of ‘Timepoint’ on 2^-ΔΔCt^, while regulating for Biological replicate.

**Hormones**

Hormone levels were quantified from leaf number 8. Five leaves were pooled at random to produce four replicates per developmental stage. After harvest, leaves were flash-frozen in liquid nitrogen and stored at -80^o^C. The samples were ground twice for 0.5 min at 30Hz with a Retsch Mixermill and were cooled with liquid nitrogen before and between milling steps. They were then extracted in 2 x 750 µl (total 1500 µl) EtAc with 0.1% formic acid, containing the internal standard (3HOBA 60 ng, DHJA 80 ng and 5IFA 50 ng per ml), for 10 min in an ultra-sonic bath. After centrifugation, the supernatant was used to quantify soluble hormones, whereas the pellet was hydrolyzed to release any bound compounds. After removal of the solvent from the supernatant by using an Eppendorf Vacuum concentrator (Mode HV) at 30 mbar, 70 µl of a fresh 1:1 mix of MeOH and TMSDM (Trimethylsilyldiazomethane, Aldrich) was added to the dry samples. GCMS (Shimadzu TQ8040) in the splitless MRM Mode was used for sample analyses. Hydrolysis of the dry pellet was performed using 200 µL of 3M HCl and 200 µL of 3M NH_3_.

**Hydrogen peroxide**

H_2_O_2_ was quantified from 20 leaves per developmental stage exactly as described in Bresson et al., (2018). A stock solution was prepared containing 0.4 mg 5(6)-Carboxy-H2DCFDA solved in 400 μL DMSO, diluted 1:1 with distilled water. The working solution contained 400 μL of stock solution added to 39.6 mL MS-Medium. Individual leaves were weighed in and incubated in 1ml working solution for 45 min at room temperature. The sample leaf was rinsed with distilled water and frozen in liquid nitrogen. An aliquot of 500 µl of 40mM Tris buffer, pH 7, was added and the sample was homogenized. After centrifugation for 15 mins at 14000 g and 4^o^C, the supernatant was measured on a plate reader (Excitation: ~480 nm, Emission: ~520 nm). On each sampling day, calibration was performed as follows. An aliquot of 0.755 mL of 0.5 M NaOH solution was mixed with 1 mL working solution. After incubation for 30 minutes at room temperature, 500 μL of 30 % H_2_O_2_ solution was added to 156 μL deacetylated dye. The sample was shaken at room temperature for 1 hour in the dark (holes were punched in tube cap to allow gas evaporation). After the addition of 226 μl Tris buffer, the fully oxidised deacetylated dye was measured photometrically at 520 nm.

**WGBS sequence processing and analysis**

The methylome raw dataset is available upon publication in the Gene expression omnibus (GEO) database under the accession number XXXXXXXX. The quality of the raw reads was assessed using FastQC Version 0.11.9. Trimmomatic 0.39 was used for the trimming and filtering of the raw reads at the following settings: SLIDINGWINDOW:4:15 LEADING:3 TRAILING:3 ILLUMINACLIP:2:30:10:1 MINLEN:36 HEADCROP:15. Sequence alignment was performed using Bismark 0.22.3 at default settings (Krueger and Andrews, 2011). The reference genome was TAIR10 assembly. Cytosine methylation was called with the bismark_methylation_extractor at the following settings: -p --comprehensive --bedGraph --CX --multicore 3 --cytosine_report. After analysis of coverage distribution, cytosines with a coverage of <3 or >40 were removed from further analyses by using R. Statistical analysis of differential methylation was carried out using the DSS package from bioconductor (Wu et al., 2013; Feng et al., 2014; Wu et al., 2015; Park and Wu, 2016). All 12 samples were combined using the *makeBSseqData* function. Data transformation and analysis were performed with *DMLtest*. For general descriptive analyses, the mean value for each position was taken from the resulting tables following all possible pairwise comparisons of the four time-points under investigation. Differential methylated loci (DML; 25 % chage) were called using the *callDML* function with *p.threshold* = 0.05. Differential methylated regions were called using the *callDMR* function with *p.threshold* = 0.05.

PCA and cluster analysis were carried out on the weighted means for the methylation of 50 bp bins of the entire genome and of all single cytosines recognized as DMLs in any of the possible pairwise comparisons. The R function *prcomp(center = T)* was used for PCA analysis followed by *ggbiplot* for plotting. Cluster analysis was performed with the R function *dist(method = ‘manhattan’)* followed by *hclust(method = ‘ward.D’)* and plotted via *plot().*

The locations of transposon elements (TEs) and gene open reading frames (ORFs) were downloaded from the TAIR10 assembly and customized into *bed* file format by using python. The region lying 2500 bp upstream of the ATG start codon of every ORF was also recorded in *bed* format and analysed as the 2500-bp promoter. The *bedtools* functions *intersect* and *closest* were used to find the locations of all DML and DMRs with respect to the above-mentioned genomic elements.

The locations of all W-boxes (TTGACT/C) were sought in the TAIR10 assembly with the regular expression function of python, namely *re.finditer*. The start and end position, together with the chromosome number, was recorded in a *bed* file format and further processed with bedtools *intersect* and *closest*. Bootstrap analysis was performed to estimate the statistical significance of the cytosine methylation in W-boxes compared with their genomic context. The average cytosine methylation level for all CHH and CHG in ORFs, TEs and 2500bp promoters was calculated separately. These were then compared with the sub-samples comprising all CHH and CHGs present in W-boxes. 10,000 random samples were drawn without replacement, with the same size as the sub-samples, and the percentage of them with the same or more extreme average was taken as the *p-*value estimate.

Differentially methylated regions (DMRs) in the *ddc* mutant were quantified from the data of Stroud et al. (2013) according to Jühling et al. (2016). Metilene 0.23 was used at the default setting and the minimum distance between the adjacent cytosines was set at 100 bp. DMRs were intersected with the locations of known genomic regions in the face of transposable elements (TEs), open reading frames (ORFs) and the -2500bp regions from the ATG start codon of ORFs (2500bp promoters) via the bedtools command *intersect*. The Softberry Nsite-PL (Solovyev et al., 2010; Shahmuradov and Solovyev, 2015) algorithm was employed to locate known transcription factor binding motifs within DMRs intersecting with 2500bp promoters. All transcription factor binding motifs specific for *A.thaliana* were then used for motif pattern recognition via MEME (Bailey et al., 2009).

**Protein expression**

WRKY18, WRKY25 and WRKY53 were expressed in *E. coli* with an N-terminal-fused 6xHis-tag, as described by Doll et al., (2020). The *E. coli* were grown overnight in 5 ml selective medium (5 ml LB + 5 µl Ampicillin 100 µg ml^-1^). An aliquot of 50 ml of the same selective medium was then inoculated with the pre-culture and shaken at 200 rpm and 37^o^C for 1.5 h. At an OD600 > 0.6, the culture was induced with 50 µl 1M IPGT and incubated overnight (18^o^C, 180 rpm). Cells were collected (30 min centrifugation at 4600 rpm, 4^o^C) and suspended in protein extraction buffer (200 mL containing 10 mM HEPES, 50 mL 1M KCl, 40.8 mL of 98% Glycerol). After sonification, the solution was centrifuged at 4^o^C and 12,000 rpm. The Bradford assay (Bradford, 1976) was used to detect protein concentrations in the crude extract.

**Verification via Western blot**

The presence of 6xHis-tagged protein in the crude extract was validated via Western blot analysis. For the SDS-PAGE, 12.8 ml water, 10.7 ml of 30% acrylamide, 8 ml of 1.5 M Tris pH 8.8 and 320 µl of 10% SDS were added together. An aliquot of 320 µl of 10% APS was mixed briefly with 32 µl TEMED. The gel was then poured into frames that were subsequently sealed with 1% agarose solution. After the acrylamide had polymerized, the gel was packed and left at 4^o^C until needed. SDS-PAGE separation was performed with 25 µg raw protein extract for approximately 2h at 330A. Semi-dry transfer to a binding membrane was performed using a transfer buffer (192mM Glycine, 25mM Tris, 200 ml of 100% EtOH) and incubation at 300 mA for 1 h.

Blocking solutions were prepared comprising 3% and 1.5% solutions of milk powder in TBS-T buffer. The membrane was first incubated in the 1.5% solution at 4^o^C for 30 mins and then blocked with 3% milk powder in TBS-T for 1 h. After 3 washing steps in TBS-T buffer for 10 mins each, the membrane was covered in a 1:5000 dilution of antibodies (anti-His-HRP conjugated antibodies, Qiagen) and incubated for 60 mins with slow rotation, followed by 3 more rounds of washing and, finally, detection.

**DPI-ELISA**

DPI-ELISA was performed following the instructions of Brand et al. (2010). A 29-bp long biotinylated artificial sequence, containing 3 W-boxes, was used for the assay:

Art. 3*Wbox F - Biotin-GGAACTTGACCATTTGACCTTGACCATGG

Art. 3*Wbox R - CCATGGTCAAGGTCAAATGGTCAAGTTCC

The sense and antisense oligonucleotides were diluted (2 pmol µl^-1^), heated for 5-10 min at 95^o^C and left to cool. Streptavidin-coated pre-blocked clear 96-well plates (Thermo Scientific Nunc Immobilizer Streptavidin, Thermo Fisher Scientific) were used for the assay. The wells were pre-washed 3 × 200 μl/well with 5xSSCT buffer (750 mM NaCl, 75 mM Sodium citrate, 0.05% (v/v) Tween20). Two concentrations of primers were tested: 0.2 and 0.02, corresponding to 20 and 2 pmol 60 µl^-1^. The annealed primer pairs were diluted to their final concentration in 5xSSCT and added to the streptavidin-coated plate, which was then incubated at room temperature for 1 h (60 µl well^-1^). The wells were washed in 2 × 100 μl 2xSSCT (300 mM NaCl, 30 mM Sodium citrate, 0.05% (v/v) Tween20) and 2 × 100 μl protein dilution buffer. Up to five crude protein extract concentrations were tested on the immobilized ds-bio DNA (0.5 μg, 5 μg, 25 μg, 50 μg, 100 μg in 60 μl per well). After being incubated for 1 h at room temperature, the plates were washed three times in 100 μl TBS-T. α-His-HRP antibody conjugate (Qiagen) diluted 1:1000 in TBS-T was added to the wells and incubated for 1h at room temperature (60 μl per well) with mild agitation. After being washed in 2 × 100 μl TBS-T and 2 × 100 μl TBS, the plates were prepared for photometric detection by being incubated with OPD solution (60 μl well^-1^ comprising 4 mg *ortho*-phenylenediamine (OPD-tablets from Sigma), 3 μl 30% H_2_O_2_ in 6 ml CP-buffer (10 mM Na_2_HPO_4_, 100 mM citric acid, pH 5 with NaOH)) in the dark for a maximum of 30 minutes. After the addition of an equal volume of stopping solution (60 μl 2N HCl), the plate was kept for a further 10 minutes in the dark. Absorbance was measured at 492 nm by using 650 nm (plate background) as a reference wavelength in an ELISA reader.

**Cytokinins**

For the estimation of cytokinin levels in the early stages of leaf senescence, Col-0 plants were grown in soil under short days for 20 days, after which flowering was induced via transition to long days. A first sample was taken at 20DAS, as an estimation for cytokinin levels before flower induction (Pre-F). Second and third samples corresponded to the Bolting (Blt) and Flowering (Flwr) stages of the previous experiment. Leaf number 8 was harvested from 12 plants per sampling point. The leaves were pooled into 3 biological replicates. The levels of t*rans*-Zeatin, *trans*-Zeatinriboside, *cis*-Zeatinriboside and Kinetin were estimated as follows. As an internal standard, 20 ng [2H5]-trans-Zeatin was added to the sample and incubated for 15 min. Extraction was performed with 1.5 ml of 0.1% formic acid and 600 mg ceramic beads in a FastPrep (6.5 M/s for 45 min) followed by incubation in a rotating vortex (99 rpm and shaking) at room temperature for 30 min. Samples were centrifuged at 12°C at 16,000 g for 30 min. The supernatant was collected and extraction was repeated with 1.2 ml formic acid (0.1%). SPE columns (Waters Oasis MCX, 150 mg) were conditioned with 5 ml methanol followed by 5 ml of 0.5 M formic acid before the sample was applied. Samples were washed first with 3 ml of 0.5 M formic acid and then with 0.1% formic acid/MeOH (95%/5% v/v). Cytokines were eluted with 3 ml NH_4_OH/MeOH (1%/60% v/v) and elutions were dried under a nitrogen flow in an ExcelVap at 22 °C. Pellets were resuspended in 150 µl of 0.1% formic acid/MeOH (95%/5% v/v). Aliquots of 15 µl were analysed by LC-MS.

**References**

Bailey TL, Boden M, Buske FA, Frith M, Grant CE, Clementi L, ... Noble WS (2009) MEME SUITE: tools for motif discovery and searching. Nucl Acid Res 37: W202-W208

Bradford MM (1976) A rapid and sensitive method for the quantitation of microgram quantities of protein utilizing the principle of protein-dye binding. Anal Biochem 72, 248–254

Brand LH, Kirchler T, Hummel S, Chaban C, Wanke D (2010) DPI-ELISA: a fast and versatile method to specify the binding of plant transcription factors to DNA in vitro. Plant methods 6: 25

Bresson, J., Bieker, S., Riester, L., Doll, J., Zentgraf, U. (2018). A guideline for leaf senescence analyses: from quantification to physiological and molecular investigations. J Exp Bot 69: 769-786

Condon DE, Tran PV, Lien YC, Schug J, Georgieff MK, Simmons RA, Won KJ (2018) Defiant:(DMRs: easy, fast, identification and ANnoTation) identifies differentially Methylated regions from iron-deficient rat hippocampus. BMC bioinformatics 19: 31

Doll J, Muth M, Riester L, Nebel S, Bresson J, Lee HC, Zentgraf U (2020) Arabidopsis thaliana WRKY25 transcription factor mediates oxidative stress tolerance and regulates senescence in a redox-dependent manner. Frontiers in Plant Science, 10: 1734

Feng H, Conneely K, Wu H (2014) A bayesian hierarchical model to detect differentially methylated loci from single nucleotide resolution sequencing data. Nucl Acid Res 42: e69

Jühling F, Kretzmer H, Bernhart SH, Otto C, Stadler PF, Hoffmann S (2016) Metilene: fast and sensitive calling of differentially methylated regions from bisulfite sequencing data. Genome Res 26: 256-262

Krueger F, Andrews SR (2011) Bismark: a flexible aligner and methylation caller for Bisulfite-Seq applications. Bioinformatics 27: 1571-1572

Park Y, Wu H (2016) Differential methylation analysis for BS-seq data under general experimental design. Bioinformatics 32: 1446-1453

Shahmuradov I. Solovyev V (2015) Nsite, NsiteH and NsiteM computer tools for studying transcription regulatory elements. Bioinformatics 31: 3544-3545

Solovyev VV, Shahmuradov IA, Salamov AA (2010) Identification of promoter regions and regulatory sites. Methods Mol Biol 674: 57-83

Wu H, Wang C, Wu Z (2013) A new shrinkage estimator for dispersion improves differential expression detection in RNA-seq data. Biostatistics 14(2):232-43

Wu H, Xu T, Feng H, Chen L, Li B, Yao B, Qin Z, Jin P, Conneely KN (2015) Detection of differentially methylated regions from whole-genome bisulfite sequencing data without replicates. Nucl Acid Res 43: e141
